# Dimerisation of European robin cryptochrome 4a

**DOI:** 10.1101/2023.04.11.536350

**Authors:** Maja Hanić, Lewis M. Antill, Angela S. Gehrckens, Jessica Schmidt, Katharina Görtemaker, Rabea Bartölke, Tarick J. El-Baba, Jingjing Xu, Karl W. Koch, Henrik Mouritsen, Justin L. P. Benesch, P. J. Hore, Ilia A. Solov’yov

**Affiliations:** Institute of Physics, Carl von Ossietzky University of Oldenburg, Carl-von-Ossietzky Straße 9-11, 26129, Oldenburg, Germany; Graduate School of Science and Engineering, Saitama University, 255 Shimo-okubo, Sakura Ward, Saitama 338-8570, Japan; JST, PRESTO, 4-1-8 Honcho, Kawaguchi, Saitama 332-0012, Japan; Department of Chemistry, University of Oxford, Physical & Theoretical Chemistry Laboratory, South Parks Road, Oxford OX1 3QZ, UK; Department of Biology and Environmental Sciences, Carl von Ossietzky University of Oldenburg, Carl-von-Ossietzky Straße 9-11, 26129, Oldenburg, Germany; Department of Neuroscience, Division of Biochemistry, Carl von Ossietzky University of Oldenburg, D-26111, Oldenburg, Germany; Kavli Institute for NanoScience Discovery, Dorothy Crowfoot Hodgkin Building, University of Oxford, OX1 3QU; Research Center for Neurosensory Sciences, Carl von Ossietzky University of Oldenburg, Carl-von-Ossietzky Straße 9-11, 26111, Oldenburg, Germany; Center for Nanoscale Dynamics (CENAD), Carl von Ossietzky Universität Oldenburg, Ammerländer Heerstr. 114-118, 26129 Oldenburg, Germany

## Abstract

Homo-dimer formation is important for the function of many proteins. Although dimeric forms of cryptochromes (Cry) have been found by crystallography and were recently observed *in vitro* for European robin Cry4a, little is known about the dimerisation of avian cryptochromes and the role it could play in the mechanism of magnetic sensing in migratory birds. Here we present a combined experimental and computational investigation of the dimerisation of robin Cry4a resulting from covalent and non-covalent interactions. Experimental studies using native mass spectrometry, mass spectrometric analysis of disulphide bonds, chemical cross-linking and photometric measurements show that disulphide-linked dimers are routinely formed, the most likely cysteines being C317 and C412. Computational modelling and molecular dynamics simulations were used to generate and assess a number of possible dimer structures. The relevance of these findings to the proposed role of Cry4a in avian magnetoreception is discussed.

## Introduction

Oligomerisation of proteins can change their structural stability, activity, and mechanisms of action^1,2^ and is important in numerous processes in living organisms. Examples from the cryptochrome (Cry) protein family include a tetrameric structure of *Arabidopsis thaliana* Cry2 (*At*Cry2) important in photoactivation^3^, a proposed dimer of L-Cry involved in the reproduction cycle of the bristle worm, *Platynereis dumerilii*^4^, and dimerisation of *At*Cry1^5, 6^.

In the investigation reported here, we focus on Cry, a protein found in all biological kingdoms^7–12^. Crys are structurally similar to light-activated DNA photolyases^7, 12, 13^ and consist of two main structural regions: a highly conserved N-terminal photolyase homology region (PHR) and the intrinsically disordered C-terminal tail (CTT)^14^. Most Crys have a flavin adenine dinucleotide cofactor (FAD) that absorbs UV-blue light, non-covalently bound in the PHR domain^7^. Crys can exist in the so-called dark-state, containing the fully oxidised form of the FAD, or in higher energy states formed by photo-excitation of the FAD followed by the formation of radical pairs, involving a triad or tetrad of tryptophan residues, which have been proposed as mediators of the mechanism of magnetoreception in migratory songbirds^15–, 20^. Three different Cry-genes, Cry1^21–26^, Cry2^21, 27^, and Cry4^28–32^ exist in most bird species, each with at least one splice-variant^22, 33, 34^. Several of these cryptochromes are found in the birds’ retina, where they can easily be light-activated^21, 24, 25, 27, 29, 32, 34^. Light-activated Cry is believed to form a signalling state, via a conformational change in the CTT, which has been the subject of computational studies of European robin Cry4a (*Erithacus rubecula*, *Er*Cry4a)^35^ and pigeon Cry4a (*Columba livia*, *Cl*Cry4a)^36^ and experimental investigations of Crys from the fruit fly (*Drosophila melanogaster*, *Dm*Cry)^37^, chicken (*Gallus gallus*, *Gg*Cry4a)^38^ and *Arabidopsis* (*At*Cry1)^39^. The exact process by which Cry combines photo-excitation with detection of the Earth’s magnetic field to form a long-lived signalling state is unknown but could conceivably involve dimerisation. The importance of homo-oligomerisation of plant Crys *in vivo* is clear^40, 41^ and optogenetic studies of *At*Cry2 show that the full-length protein undergoes light-induced oligomerisation and that functionality is lost in a truncated form that lacks the CTT domain^42^.

For a signal transduction cascade to function, Cry must interact with other proteins. Wu *et al.* recently listed six proteins as candidate *Er*Cry4a interaction partners^43^. A subsequent detailed biochemical investigation of *Er*Cry4a and a cone-specific G-protein from European robin demonstrated that these two proteins interact directly with each other^44^. Dimers of *Er*Cry4a have also been reported^15^, but is unclear whether/how a monomer-dimer equilibrium might be involved in the downstream signalling process. Within the Cry family there are reports of crystallographic dimeric asymmetric units, including *Dm*Cry^45^ and mouse (*Mus musculus*) *Mm*Cry1 and *Mm*Cry2^46^ (Fig. 1). A study of full-length *Dm*Cry noted an intermonomer disulphide bond, but deemed it to be an artefact of the crystallisation method^47^. To date, there has been little work on the dimers of animal Crys, and their function, if any, is unknown.

**Figure 1.**
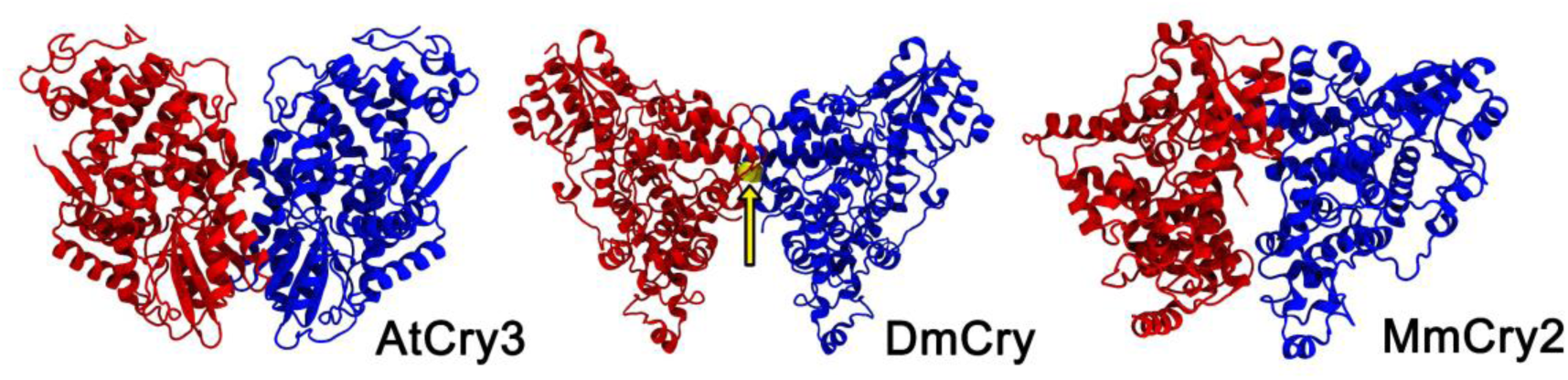
Structures of cryptochrome dimers from the Protein Data Bank (PDB): non-covalent *At*Cry3 dimer (PDB: 2J4D^3^), covalent *Dm*Cry dimer (PDB: 4GU5^28^) (the arrow indicates the C296-C296 disulphide bond) and non-covalent *Mm*Cry2 dimer (PDB: 6KX8^29^).

To learn more about *Er*Cry4a dimerisation and its possible role in magnetoreception^15, 18^, we have explored a variety of candidate structures, including covalently and non-covalently linked forms of the full-length and truncated protein, using a combination of experimental and computational methods to identify potential *Er*Cry4a dimers. Native mass spectrometry (MS), mass photometry (MP), and gel electrophoresis of wild-type (WT) and mutant proteins were used to establish the presence and nature of the dimers, while chemical cross-linking followed by mass spectrometry (XL-MS) provided information about the relative orientation of the monomer units. A combination of molecular docking and molecular dynamics (MD) techniques provided model structures for comparison with the experimental data.

## Methods

### Experimental Methods

#### Protein expression and purification

WT *Er*Cry4a (GenBank: KX890129.1) was cloned, expressed and purified as described by Xu *et al.*^15^ with the following modifications. The LB media contained 10 g L^−1^ yeast extract and the expression time was 44 h instead of 22 h. The mutants were generated using the Q5 site-directed mutagenesis kit (New England Biolabs, Ipswich, MA, USA), and plasmids were confirmed by Sanger sequencing (LGC Genomics). For the C412A mutant and the 5-cysteine mutant (C68A+C73A+C116A+C189A+C317A), an *E. coli* K12 codon-optimised version of *Er*Cry4a was used, generated by Eurofins Genomics (Ebersberg, Germany). Tables 1 and 2 summarise the primers used for each mutation. Mutant proteins were expressed and purified in the same way as the WT protein with the following exception: double, triple and 5-cysteine mutants were expressed in BL21-CodonPlus(DE3)-RIPL *E. coli* cells (Agilent, Santa Clara, CA, USA) in the dark, starting with a 30 mL culture grown overnight at 30 °C and 250 rpm, a preculture inoculated to an OD_600_ of 0.05 for 6–6.5 h at 37 °C, and a main culture inoculated with the preculture to an OD_600_ of 0.5 for about 3 h at 37 °C until the OD_600_ reached 0.6, when the shakers were set to 15 °C and 160 rpm. After about 45 min, at an OD_600_ of 0.9 to 1.0, protein expression was induced with 5 µM isopropyl β-D-1-thiogalactopyranoside. Cell harvest, lysis and purification using Ni-NTA agarose columns and anion exchange chromatography were all performed as described in Xu *et al.*^15^ Purified protein samples were either used directly for photometric cysteine exposure measurements or buffer-exchanged, to remove reducing agents, into 20 mM Tris, 250 mM NaCl, and 20% glycerol using Sephadex G 25 in PD10 desalting columns (Cytiva, Uppsala, Sweden). Protein samples for XL-MS were produced in insect cells essentially as described previously^44^ with the additional step of removing the His-tag. To cleave the His-tag from the Ni NTA eluted protein, samples were buffer-exchanged into 20 mM Tris, 100 mM NaCl, 2 mM *DL*-dithiothreitol, DTT, using Sephadex G 25 in PD10 desalting columns (Cytiva) and the His-tag was cut off overnight at 4 °C by the addition of 10 U AcTEV Protease (Thermo Fisher). *Er*Cry4a was further purified using anion-exchange chromatography as previously described^44^. All purified protein samples were concentrated to ∼3.5 mg mL^−1^. Samples were snap-frozen in liquid nitrogen and stored at −80 °C until shipment to Oxford on dry ice.

**Table 1.**
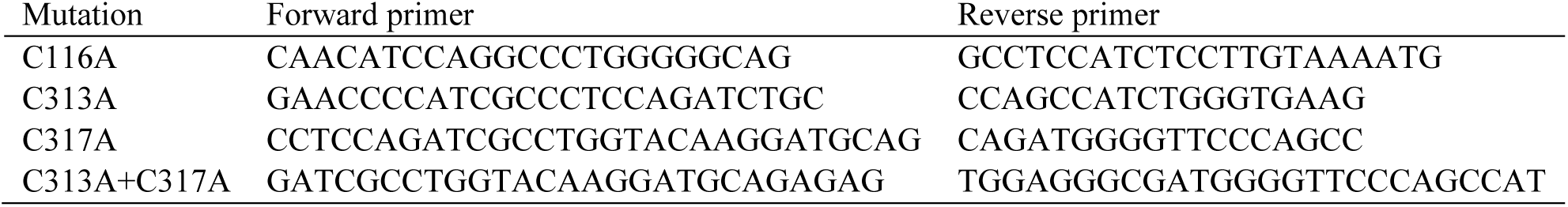
Mutations introduced in the non-codon-optimised ErCry4a.

**Table 2.**
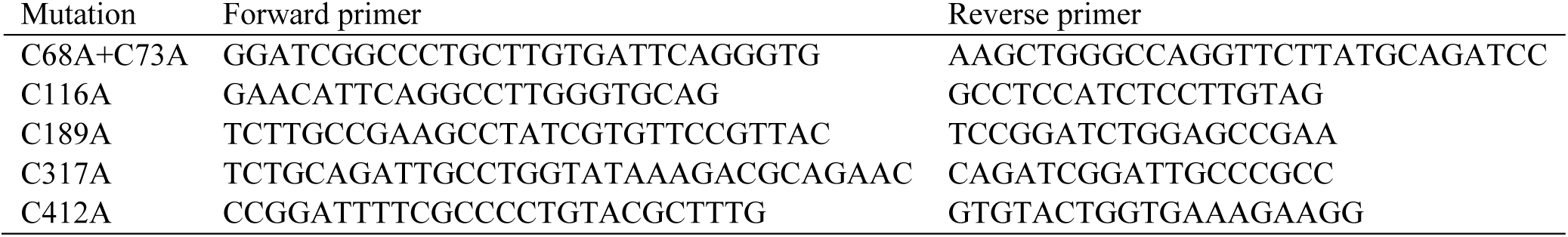
Mutations introduced in the codon-optimised ErCry4a.

#### Native Mass Spectrometry and SDS-PAGE

Native MS and sodium dodecyl sulphate–polyacrylamide gel electrophoresis (SDS-PAGE) were used to investigate the dimeric nature of *Er*Cry4a. The *Er*Cry4a samples used for these experiments were shipped overnight to Oxford on dry ice from the production laboratory at the University of Oldenburg, in a buffer containing 20 mM Tris, 250 mM NaCl, and 20% glycerol at pH 8. The samples were kept at −80 °C until they were thawed on ice and buffer-exchanged into 200 mM ammonium acetate, pH 8, for the native MS measurements using Zeba™ Micro Spin desalting columns with a molecular weight exclusion limit of 40 kDa (ThermoFisher Scientific). All samples included an N-terminal 10×His-tag used for purification of the proteins.

Measurements were performed in Oxford using in-house gold-plated capillaries on a Q Exactive™ mass spectrometer in positive ion mode. The samples were sprayed using nanoflow electrospray with a source temperature of 150 °C and a capillary voltage of 1.0 kV^48^. The higher-energy C-trap dissociation (HCD) cell voltage was 5 V, in-source trapping was set to −200 V to help with the dissociation of small ion adducts, and the noise threshold parameter was 3. Ion transfer optics and voltage gradients throughout the instruments were adjusted for optimum protein transmission.

For SDS-PAGE gels, the samples were covered in the dark for 20 min at room temperature at a concentration of ∼8 µM. 3.2 µg of each sample were then mixed with diluted NuPAGE LDS sample buffer (4X) (Invitrogen, Waltham, MA, USA) before being loaded on NuPAGE Novex 4-12% bis-Tris gels, 1.0 mm × 10 well (Invitrogen) and run with NuPAGE MES SDS running buffer (20X) (Invitrogen). Samples of proteins without a His-tag, containing 2 mM DTT (Sigma-Aldrich) were shipped to Oxford and kept in the light while being incubated with different concentrations of DTT for 20 min at room temperature prior to being submitted to the SDS-PAGE gel. Protein bands were visualised by Coomassie blue staining using QuickBlue Protein Stain (LuBio, Zürich, Switzerland) and accompanied by either a SeeBlue^TM^ Plus2 Pre-stained Protein Ladder (Invitrogen) or a PageRuler Prestained Protein Ladder (Thermo Scientific, Waltham, MA, USA). The software GelAnalyzer 19.1^49^ was used for evaluation of the gel bands.

#### Mass photometry

All samples were shipped as described above except for the *Er*Cry4a WT sample used in Fig. 3 which was shipped with the addition of 2 mM DTT to limit higher order oligomerisation during shipping. This sample was diluted to a concentration of ∼44 nM in 200 mM ammonium acetate, pH 8.3, prior to measurement. All other samples were buffer-exchanged into 200 mM ammonium acetate, pH 8.0, and diluted to 13-15 nM. MP measurements were performed essentially as described previously (Young *et al.*^50^), on a ONE^MP^ instrument (Refeyn Ltd, Oxford) in the case of the *Er*Cry4a WT sample in Fig. 3 and on a TWO^MP^ instrument (Refeyn Ltd, Oxford) in all other cases. Briefly, a borosilicate microscope coverslip (Thorlabs) was cleaned by ultra-sonication for 5 min in a 1:1 mix of ultrapure water and isopropanol followed by another 5 min of ultra-sonication in ultrapure water and then dried under a stream of nitrogen. Cleaned coverslips and Grace Bio-Labs reusable CultureWell^TM^ gaskets (Bio-Labs) were assembled into flow chambers. 10-20 μL of sample were added to the flow chamber and images of a 3.5 × 12.2 μm^2^ region of the glass coverslip surface were acquired at 1000 frames s^-1^. The mass distributions were obtained from contrasts of ratiometric data by calibration using DiscoverMP (Refeyn Ltd, Oxford).

#### Mass spectrometric analysis of chemically cross-linked peptides

XL-MS was used to probe the interaction surfaces of the linked monomers. Samples for XL-MS were sent to Oxford as described above, with the addition of 2 mM DTT in the shipping buffer to limit higher-order oligomerisation. Furthermore, His-tags were cut leaving only 5 additional amino acids at the N-terminus (GAMGS).

To investigate potential disulphide bonds, protein solutions were incubated with the linkers disuccinimidyl sulphoxide (DSSO)^51^ or disuccinimidyl dibutyric acid (DSBU)^52^ to cross-link primary amines at the dimer interface using the manufacturer’s recommended procedure, as described previously.^53^ Cross-linked and non-cross-linked proteins were resolved using SDS-PAGE for analysis of the cross-linkers and direct analysis of the disulphide bonds.

Bands were excised from gels and prepared for mass spectrometry as described by Shevchenko *et al.*^54^ After in-gel digestion, peptides were resolubilised in buffer A (H_2_O, 0.1% formic acid), and ∼5-10 µL of peptide was loaded onto a reverse-phase trap column (Acclaim PepMap 100, 75 µm × 2 cm, NanoViper, C18, 3 µm, 100 Å, ThermoFisher, Waltham, MA, USA) using an Ultimate 3000 autosampler and washed with 40 µL of buffer A at a flow rate of 20 µL min^−1^. The trapped peptides were then separated by applying a 15 cm reverse-phase analytical column (Acclaim PepMap 100 C18, 3 µm, 150 mm × 0.075 mm) using a 105 min linear gradient from 5% to 55% buffer B (80% acetonitrile, 20% water, 0.1% formic acid) at a flow rate of 300 nL min^−1^. Long and hydrophobic peptides were washed from the column using 99% buffer B for 15 min at the same flow rate. The separated peptides were electrosprayed in the positive ion mode into an Orbitrap Eclipse Tribrid mass spectrometer (ThermoFisher) operated in data-dependent acquisition mode (3 s cycle time) using the workflow recommended by the DSSO and DSBU methods programmed by the manufacturer. Briefly, precursor scans were collected in the Orbitrap analyser at 60,000 resolving power at *m*/*z* 200 with a mass range of 375-1600 *m*/*z*. Precursors above the intensity threshold of 1.0 × 10^4^ having charge states between *z* = 4 and *z* = 8 were isolated using the quadrupole (0.5 *m*/*z* offset, 1.6 *m*/*z* isolation window) and fragmented in the linear ion trap using collision-induced dissociation (collision energy = 25%). Peptide fragments were analysed using the Orbitrap at a resolving power of 30,000 at *m*/*z* 200 with a maximum injection time of 70 ms. Fragment ions with *z* = 2 to *z* = 6 spaced by the targeted mass difference of 25.9 Da or 31.9 Da (± 10 ppm) corresponding to peptides cross-linked by DSBU or DSSO, respectively, were subjected to further sequencing in the linear ion trap operated in rapid detection mode (35% collision energy, MS1 isolation of 2.5 *m*/*z* and MS2 isolation of 2.6 *m*/*z*). Additional MS/MS scans for precursors within 10 ppm were dynamically excluded for 30 s following the initial selection. The cross-linking data were analysed using the free software tool MeroX^55^ available at www.StavroX.com.

#### Cysteine exposure measurements

Quantitative determination of thiol groups in solution was achieved essentially as described previously^56^ by recording the formation of 5-thio-2-nitrobenzoic acid (TNB) from 5,5′-dithio-bis-(2-nitrobenzoic acid) (DTNB) at a wavelength of 412 nm. Briefly, WT *Er*Cry4a, mutants C317A, C412A and the 5-cysteine mutant were dialysed overnight in 100 mM Tris/HCl at pH 8.0. Next, a fresh 12 mM DTNB solution in 100 mM Tris/HCl pH 8.0 was produced. A calibration curve with 0, 5, 10, 15, 20, 25 and 30 µM L-cysteine and 60 µM DTNB was prepared to calculate the slope of the line (see Fig. S2). After adding DTNB, the samples were incubated for 10 min at room temperature. Afterwards, the absorbance of TNB in the Cry samples was measured at 412 nm. To measure the accessibility of cysteines in the proteins, 5 µM of WT *Er*Cry4a and of the *Er*Cry4a mutants were incubated for 10 min at room temperature with 60 µM DTNB in 100 mM Tris/HCl pH 8.0. The number of accessible cysteines was calculated using the expression, *N*_Cys_=*E* / *ac*, where *E* is the total absorbance of the sample, *a* is the slope of the calibration graph and *c* is the concentration of protein in the solution. All measurements were done in ambient light conditions.

### Computational Methods

Dimer formation through both covalent and non-covalent interactions of surface residues have been investigated for both full-length and truncated *Er*Cry4a structures. All of the dimers studied were classified into families representing groups of structures according to the type of interaction, i.e., covalent (cov) or non-covalent (ncov). The covalent structures were labelled cov(*N*)*^n^*, where *N* is the sequence number of the cysteine residue in the monomer subunits that forms the disulphide bond and *n* labels a particular dimer within the family.

#### Identifying exposed residues for covalent dimer construction

Covalently linked dimers arise from the formation of disulphide bonds between cysteine residues. Figure 2 summarises the overall workflow for constructing the 16 *Er*Cry4a dimeric structures we have investigated. A previously simulated *Er*Cry4a construct, including the FAD cofactor, was used for an analysis of the solvent exposure of the 11 cysteines in *Er*Cry4a^15^. 50 snapshots, taken from a 200 ns MD simulation, were time-averaged and evaluated using GETAREA2^57^. To compare SASA values for different amino acids, the absolute solvent-accessible surface area for each residue (SASA)^58^ was divided by the maximum SASA (MSA) value for the corresponding amino acid (X) in a Gly-X-Gly tripeptide. A similar method was introduced in earlier work^59^. Gly-X-Gly tripeptides were constructed with Pep McConst^60^ software and simulated for 1 ns in a waterbox using NAMD. MSA values for the X residues were determined using GETAREA2^57^. The solvent exposure for residue, *i*, was calculated as:

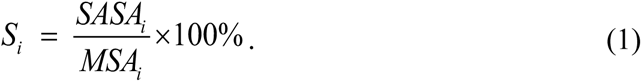

**Figure 2.**
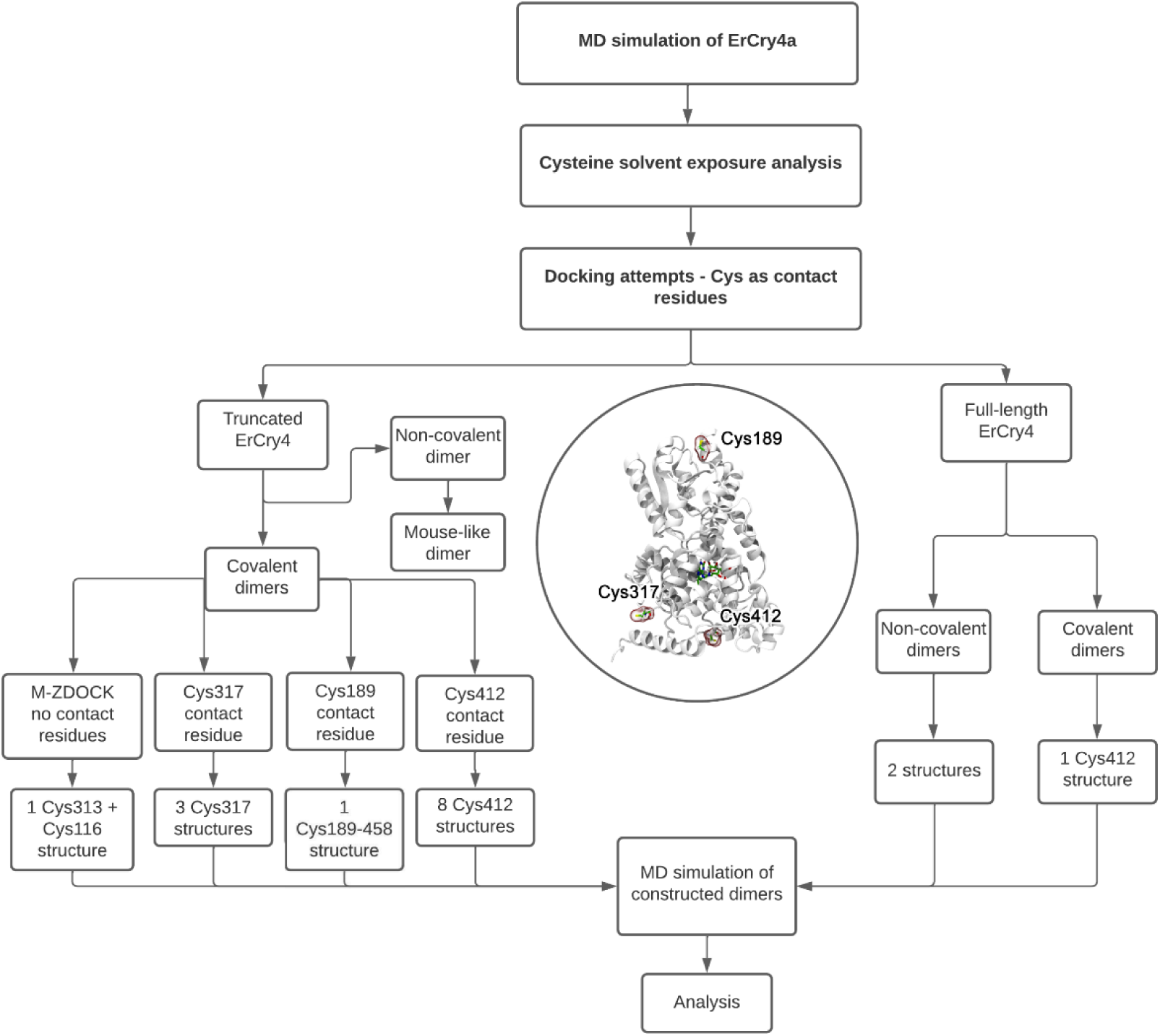
Computational workflow used to produce the 16 dimeric structures of *Er*Cry4a. The inset in the middle highlights the contact cysteine residues (C317, C412 and C189) which were used in constructing the covalent dimers. An overview of all structures obtained during the docking procedure can be found in the Supplementary Material in Tables S10-S18

#### Covalent dimer construction

The ZDOCK docking tool^61–63^ was used to create a set of *Er*Cry4a dimers using the seven most solvent-exposed cysteine residues as the docking sites (Table 3). ZDOCK allows one to define a contact residue for dimer construction such that the defined residues in the monomers are placed in close proximity to each other. For the six most solvent-exposed cysteines residues (C68, C73, C116, C189, C317 and C412) together with C179, 49 dimeric structures were constructed, and inter-cysteine distances were analysed to determine the possibility of disulphide bond formation (See Tables S10-S18). Docking of two *Er*Cry4a monomers with the contact residues Cys68 (Table S10), Cys73 (Table S11), Cys116 (Table S12), or Cys179 (Table S13) produced 10 dimeric structures. However, because the cysteine residues were more than 10 Å apart, these structures were not considered in the MD simulations (Table S19). With Cys189 (Table S14) and Cys317 (Table S15) as contact residues, four dimers were found in which one and three structures, respectively, contained the cysteines in close enough contact to warrant proceeding to MD simulations. The disulphide bonds were introduced with NAMD^64^ by removing the hydrogen atoms from the cysteine SH groups and defining an abnormally long covalent connection between the sulphur atoms. During the follow-up equilibration, the bond acquired the length of a typical disulphide bond causing a slight rearrangement of atoms at the interface to accommodate the structural change. With Cys412 as the contact residue, ZDOCK yielded three dimeric structures (Table S16) in which the cysteines were somewhat separated, making them unstable in short MD simulations. Since the experimental results (below) suggested that the Cys412 residue was potentially important for dimerisation, we have used snapshots from the short MD simulation to construct and simulate 8 dimeric configurations for the Cys412 family by first imposing an artificial constraint on the Cys412-Cys412 bond to help the residues move closer together, and then removing the constraint.

**Table 3.**
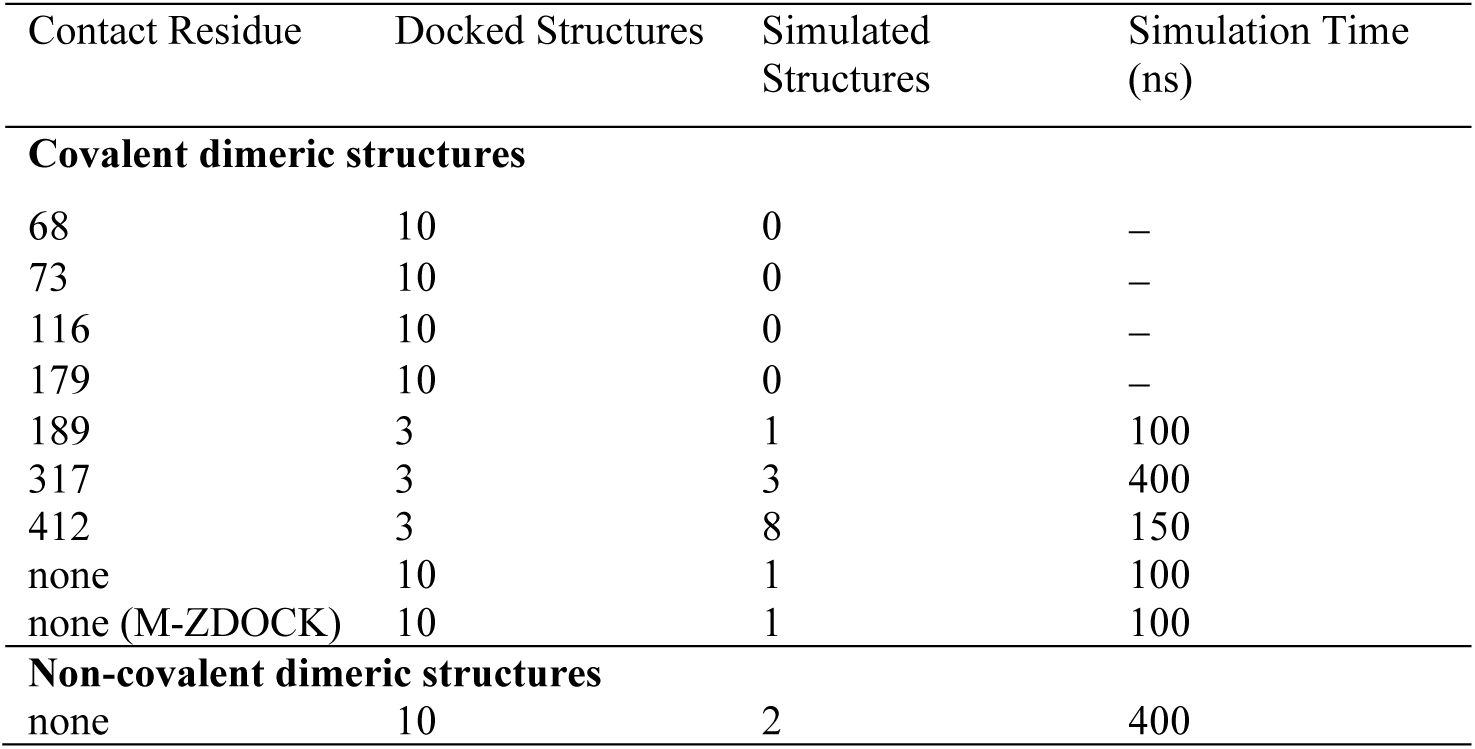
Summary of *Er*Cry4a dimer structures produced by ZDOCK and M-ZDOCK and the MD simulations. The specified lengths of the simulations apply to each member of the protein family.

| Contact Residue | Docked Structures | Simulated Structures | Simulation Time (ns) |
| --- | --- | --- | --- |
| **Covalent dimeric structures** |  |  |  |
| 68 | 10 | 0 | – |
| 73 | 10 | 0 | – |
| 116 | 10 | 0 | – |
| 179 | 10 | 0 | – |
| 189 | 3 | 1 | 100 |
| 317 | 3 | 3 | 400 |
| 412 | 3 | 8 | 150 |
| none | 10 | 1 | 100 |
| none (M-ZDOCK) | 10 | 1 | 100 |
| **Non-covalent dimeric structures** |  |  |  |
| none | 10 | 2 | 400 |

An additional 10 structures were generated without explicitly specifying a contact residue (Table S17). Furthermore, the M-ZDOCK tool^65^ was used to search for different plausible *Er*Cry4a dimer structures by predicting cyclically symmetric multimers based on the structure of the *Er*Cry4a monomer (Table S18). In the case of a dimer, ten symmetric structures were obtained, but only one, referred to as cov^D^, was analysed further (Table S18). This dimer contained two disulphide bonds: C116-C313 and C313-C116.

#### Non-covalent dimer construction

The computational tool I-TASSER^66–69^ was used to generate the intrinsically disordered CTT (residues 498-527) missing from the crystal structure of pigeon Cry4a (PDB code: 6PU0)^28^ to obtain a full-length (FL) homology model, *Er*Cry4aFL, with a (confidence) C-score of 0.34. This parameter reports the quality of the predicted model and ranges from −5 to +2, where a higher value signifies a model with a higher confidence (Table S5). The C-score is calculated based on the significance of threading template alignments with subsequent reassembly using replica-exchanged Monte Carlo simulations and the convergence parameters of the structure assembly simulations^66–69^.

Structures of possible *Er*Cry4aFL non-covalently bound dimers were generated using ZDOCK^61^. Shape complementarity was taken into consideration, where the algorithm keeps one monomer fixed and rotates and translates the other to find configurations that result in the best fitting score. In the investigation of possible covalent and non-covalent dimers, ZDOCK produced 2000 dimer configurations, of which the top 10 were used for further investigation. These 10 structures were chosen using an energy-based scoring function which includes the potential energy, spatial complementarity, and electric field force between the protein subunits.

#### Dimeric structure inspired by ***Mm***Cry2

Dimer interfaces have a high degree of conservation in evolutionarily related proteins^1^. Even though the crystallographic dimer of *Mm*Cry2 has not been identified as being functionally relevant, it is still the closest protein to *Er*Cry4a for which a dimer crystal structure has been reported (PDB: 6KX8^46^). A dimeric structure inspired by *Mm*Cry2 was thus constructed, in which the monomers were structurally aligned with the *Mm*Cry2 dimer subunits.

#### Molecular dynamics simulations

All MD simulations were conducted using the NAMD package^64, 70^ through the VIKING platform^71^. The CHARMM36 force field included CMAP corrections for proteins^72, 73^ and additional parameterisations for FAD^29, 74–76^. Periodic boundary conditions were adopted in all MD simulations and the particle mesh Ewald summation method was employed to evaluate long-range Coulomb interactions. Van der Waals interactions were treated using a smooth cut-off of 12 Å with a switching distance of 10 Å. The simulation temperature was 310 K, controlled with the Langevin thermostat^70^. A constant pressure of 1 atm for equilibrium simulations was obtained using the Langevin piston Nosé–Hoover method^77^. The SHAKE algorithm was used to constrain bonds including hydrogen atoms at their respective equilibrium distances. After 10,000 NAMD^70^ minimisation steps, harmonic restraints were initiated in the system and gradually released to achieve an equilibrium structure. Equilibration simulations were conducted with explicit solvent modelled through the TIP3P parameter set^78^ with water molecules surrounding the dimers to a distance of 15 Å in all directions. A NaCl salt concentration of 50 mM was assumed in all simulations. After the equilibration, production simulations with temperature control (310 K) within the *NVT* statistical ensembles were performed (Table S19). All MD simulation results were analysed with VMD^79^.

## Results and discussion

We have studied the dimerisation of *Er*Cry4a using experimental and computational approaches in parallel. Computational results have been used to guide the experiments and *vice versa* at various stages of the investigation.

### *Er*Cry4a forms a population of disulphide-linked dimers

Figure 3A shows a native mass spectrum of WT *Er*Cry4a. Two charge-state series consistent with the monomer and dimer can be seen: major peaks correspond to the monomer, and minor peaks in the range *m*/*z* = 5000-7000 to the dimer.

**Figure 3.**
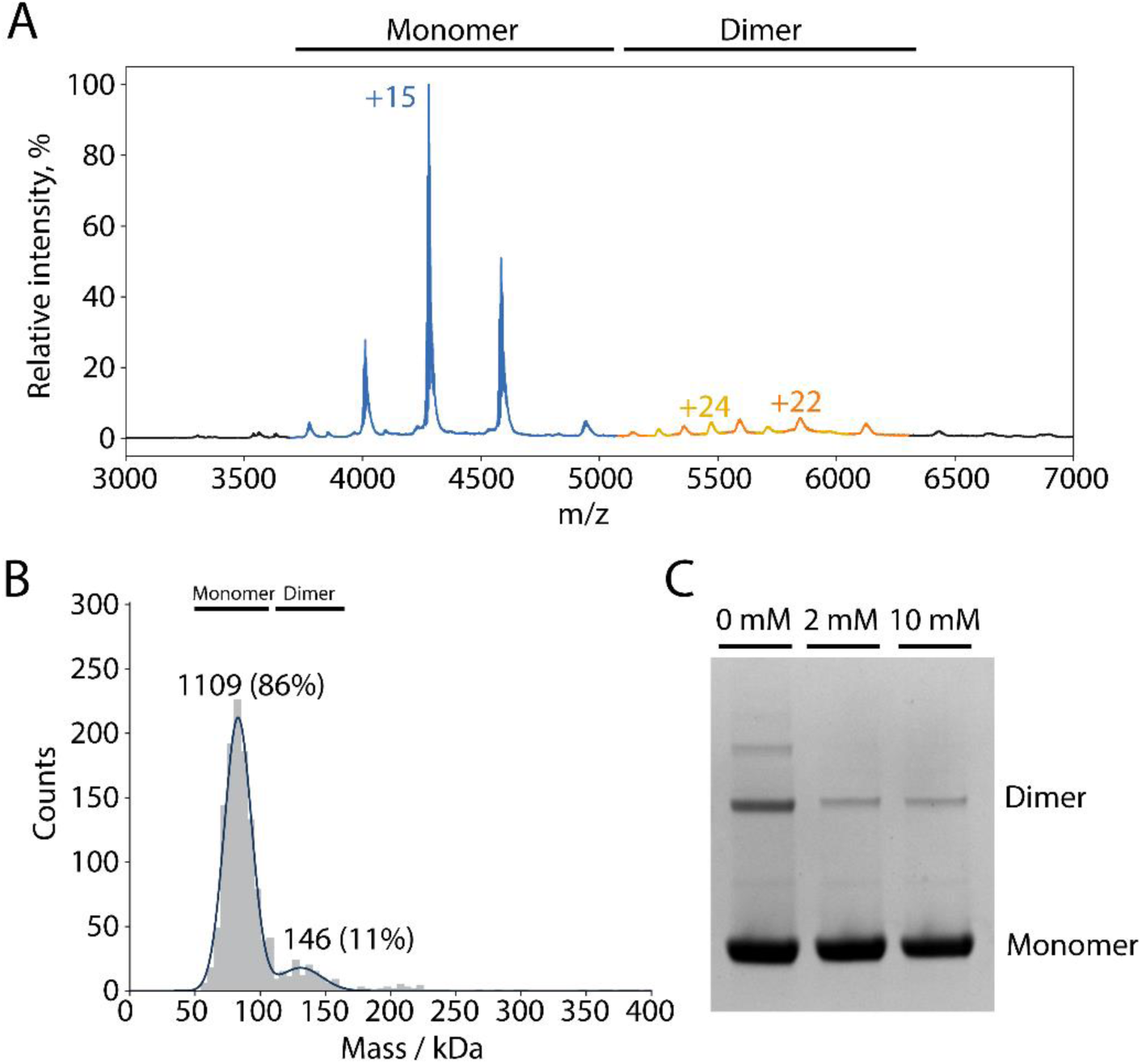
(**A**) Native mass spectrum of *Er*Cry4a. The peaks of the monomer region are indicated in blue (molecular mass 64.197 ± 0.029 kDa) and the peaks of the dimer region in orange and yellow (molecular masses ∼123 kDa and ∼131 kDa). The expected mass of an *Er*Cry4a WT monomer including FAD and a His-tag is 64.034 kDa. The differences between the detected and the expected masses are due to salt ions and buffer components. The finding of a second dimer mass, lower than the expected mass, could be due to minor truncations and is only observed in some samples. The observation here of stable dimers is consistent with Fig. S13 of Xu et al.^15^ The sample used for this measurement contained 10 mM BME during shipment to prevent higher order oligomerisation. (**B**) Mass photometry of 44 nM *Er*Cry4a. The same sample as in (A) was used. The numbers above the peaks are the numbers of counts. (**C**) SDS-PAGE gel of *Er*Cry4a samples without a His-tag (the same as used for XL-MS) that had been incubated with various amounts of DTT.

To investigate the dimerisation further, WT *Er*Cry4a was studied by mass photometry (Fig. 3B). Two peaks were observed, a major component (60-80 kDa, 86%) from monomers and a minor component (120-160 kDa, 11%) attributed to dimers. The observation of dimeric forms of *Er*Cry4a both by native MS (at micromolar protein concentrations) and MP (at nanomolar concentrations) suggests that the two monomer units are strongly bound and not an artefact of nanoelectrospray ionisation. Non-specific, non-covalent dimers would be expected to dissociate at high dilution. Notably, the relative abundance of dimers and monomers was the same in both the MS and MP experiments despite the large difference in sample concentration, suggesting that the dimers are covalently linked. To test this possibility, we ran SDS-PAGE gels of *Er*Cry4a samples after incubation with various concentrations of DTT reductant (Fig. 3C). Bands from dimers and higher order oligomers, visible under the denaturing conditions of the gels in the absence of DTT, are attenuated by DTT-treatment, consistent with dimer formation via covalent inter-monomer disulphide bonds. The complete gel is shown in Fig. S1.

### Cysteine solvent-accessibility calculations

On the basis that the covalent links responsible for dimerisation are most likely to be disulphide bonds, we calculated the solvent-accessible surface area (SASA) of the 11 cysteines in *Er*Cry4a (Fig. 4 and Table S3). Six of these residues have sidechain accessibilities greater than 20%, the other five being less than 7%. The most exposed cysteine, C317, is part of a turn near to the end of the Trp-tetrad. The next four most exposed are all located on or close to an α-helix.

**Figure 4.**
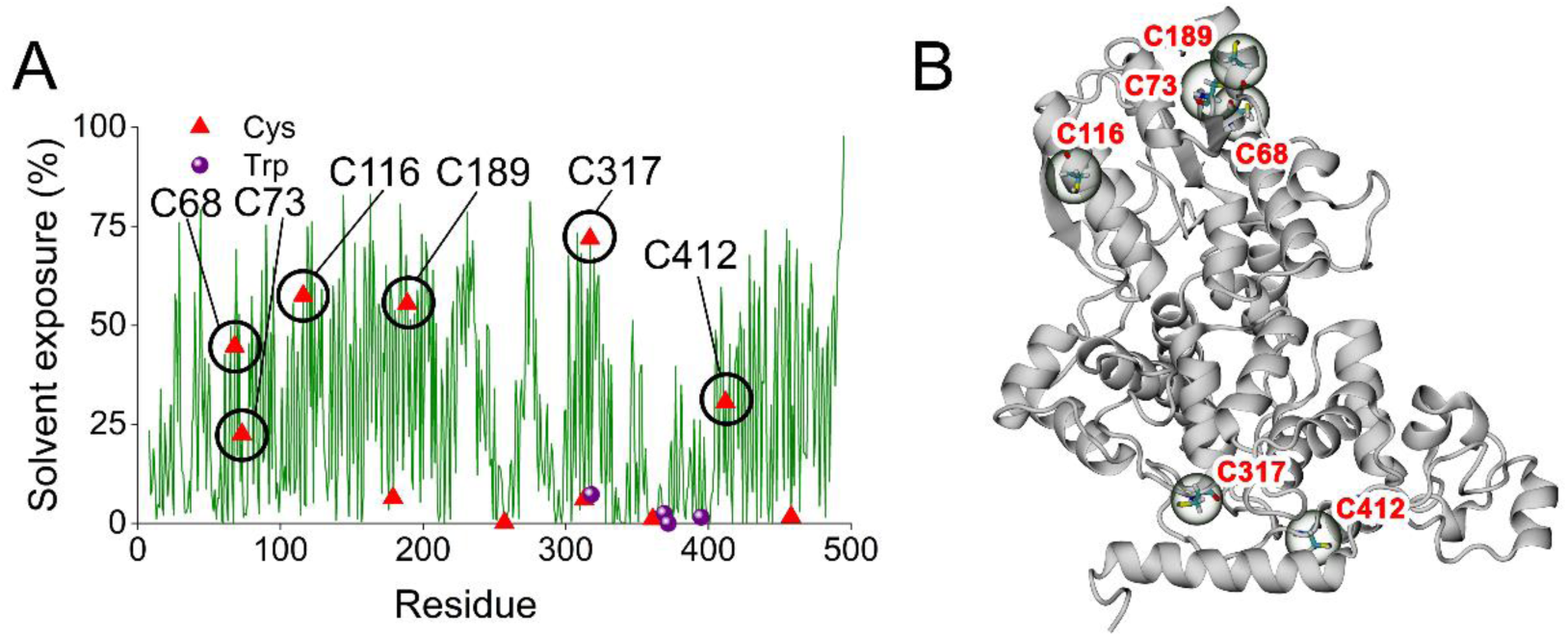
(**A**) Solvent exposures of residues in monomeric WT *Er*Cry4a with cysteines (red triangles) indicated. Also shown (purple spheres) are the components of the Trp-tetrad: (W318, W369, W372, W395). (**B**) *Er*Cry4a secondary structural elements with the six most exposed cysteines indicated.

### Dimerisation of cysteine mutants

M-ZDOCK was used to identify potential dimers by docking pairs of monomers without specifying a contact residue (see below). Using a selection criterion that the sulphur atoms of the cysteines should be closer than ∼6 Å, three residues, C116, C313, and C317, were identified as possible components of intermolecular disulphides (Table S18). C116 and C317 have the highest estimated solvent exposure of all 11 cysteines, while C313 is largely buried (Fig. 4 and Table S3). Four mutants of *Er*Cry4a were expressed and purified with one, two or all three of these cysteines replaced by alanines: C317A, C116A+C313A, C116A+C317A, and C116A+C313A+C317A. As judged by SDS-PAGE (Fig. S1B), all four mutants oligomerise, implying that if cysteines C116, C313 and C317 are involved in dimer formation, they are not the only ones.

Two further *Er*Cry4a mutants were investigated: C412A (chosen based on docking followed by MD simulations and on chemical cross-linking followed by mass spectrometric analysis, see below) and C68A+C73A+C116A+C189A+C317A (with five of the six cysteines with the largest calculated SASAs replaced by alanines). Both proteins oligomerised showing that these mutations, like the four above, might reduce the ability of *Er*Cry4a to dimerise but fail to remove it completely.

The *Er*Cry4a mutants C412A, C317A, and C412A+C317A were subsequently chosen for further comparative investigation. As seen in Fig. 5, all three have a lower degree of dimerisation than the WT protein, and the double mutant dimerises less than either of the single mutants in both MP and on the gels. In the MP measurements (Fig. 5B), the degree of dimer formation is in the order WT > C317A > C412A > C412A+C317A. The low concentrations used in this experiment (13-15 nM) suggest that this dimerisation is largely covalent and involves both cysteines. The reversed order of C317A and C412A in Fig. 5C perhaps indicates a different tendency for non-covalent and covalent association. The presence of trimers in the denaturing gel necessitates that two or more disulphide links are required for their formation.

**Figure 5.**
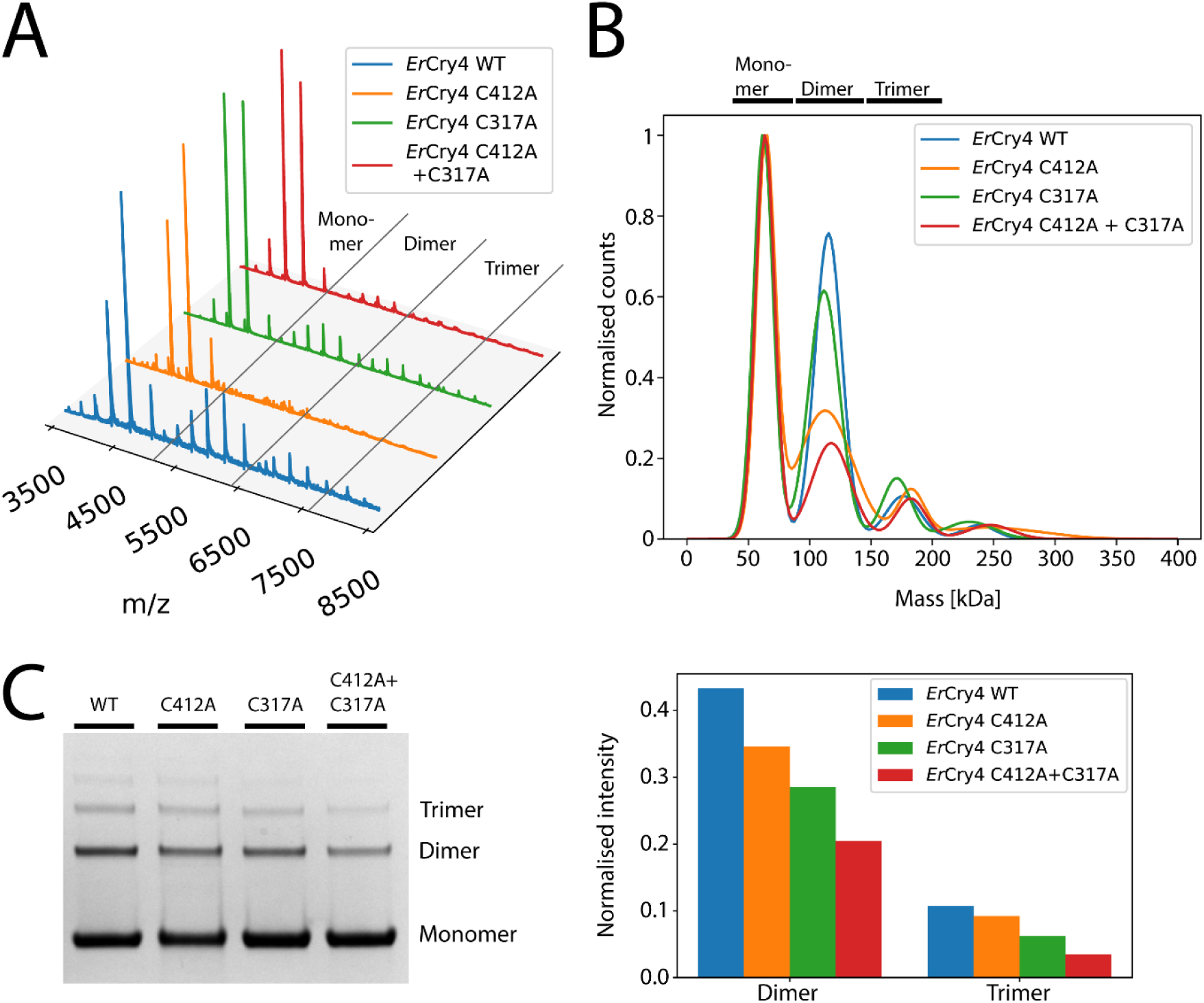
(**A**) Native mass spectra of WT *Er*Cry4a and three cysteine mutants. All four proteins are mostly monomeric but also show charge-state envelopes from dimers and trimers. (**B**) Mass photometry results for the same proteins at concentrations between 13-15 nM displayed as Gaussian-fitted normalised counts. The four traces have been scaled so that the monomer peaks have the same height. (**C**) Denaturing SDS-PAGE gel for the same four proteins (full SDS-PAGE gels can be found in Fig. S1C) and the normalised intensities of the protein areas.

### Cysteine accessibility measurements

The average number of accessible cysteines in WT and mutant forms of *Er*Cry4a was determined by measuring the absorption of the product formed by treating the protein with Ellman’s reagent (DTNB). The assay finds approximately five accessible cysteines for the wild-type protein, four for each of the C317A and C412A mutants and three for the C116A+C317A double mutant (Table 4). More detail on these absorbance measurements is given in Table S1. Assuming that each cysteine is either completely unreactive towards DTNB due to structural hindrance or reacts stoichiometrically, these results suggest that all three of C317, C412 and C116 are significantly exposed on the surface of the protein and in principle available for the formation of intermolecular disulphide bonds. However, as a note of caution, a cysteine able to react with DTNB (a small molecule) is not necessarily accessible enough for disulphide formation between monomers.

**Table 4.** TNB absorbance measurements (at 412 nm) and average number of accessible cysteines in wild-type and mutant forms of ErCry4a

| | $A_{412}$ | Accessible Cysteines |
| --- | --- | --- |
| WT | $0.30 \pm 0.01$ | $4.78 \pm 0.13$ |
| C317A | $0.27 \pm 0.01$ | $4.23 \pm 0.10$ |
| C412A | $0.27 \pm 0.00$ | $4.16 \pm 0.05$ |
| C116A+C317A | $0.17 \pm 0.01$ | $3.30 \pm 0.11$ |

### Cross-linking mass spectrometry

XL-MS was used to investigate the interface between monomer units in *Er*Cry4a dimers by creating artificial chemical cross-links that can be cleaved by MS so as to reveal which parts of the monomers come into close contact in the dimers. The bifunctional compounds DSSO and DSBU cross-link the exposed primary amine groups of lysine residues, via bis(N-hydroxysuccinimide) ester groups. MS analysis of peptides formed by enzymatic digestion of the cross-linked dimers identifies pairs of lysines with α-carbons separated by up to ∼2.7 nm thereby constraining the location and geometry of the protein-protein interface.

Five lysines were identified to form cross-links. Based on whether they were found in the monomeric or dimeric fractions of the protein, two links were assigned to be intramolecular and three to be intermolecular, as displayed in Fig. 6A. Four disulphide bonds were identified (C361-C458, C412-C361, C412-C412, and C412-C458), three of which involve C412. Of these, only C412-C412 is unequivocally intermolecular: the difficulty of distinguishing the two possibilities in a homodimer means that the other three could be intermolecular or intramolecular or both. The intermolecular K152-K152 link is consistent with a C412-C412 disulphide bond because K152 and C412 are on the same side of the protein (Fig. 6B). The links from K234 to K429 and from K152 to K507 were also found in the monomer protein fraction suggesting that they are more likely to be intra-monomer. For more details see Table S2. Overall, these results point to C412 as a likely component of disulphide links between monomers.

**Figure 6.**
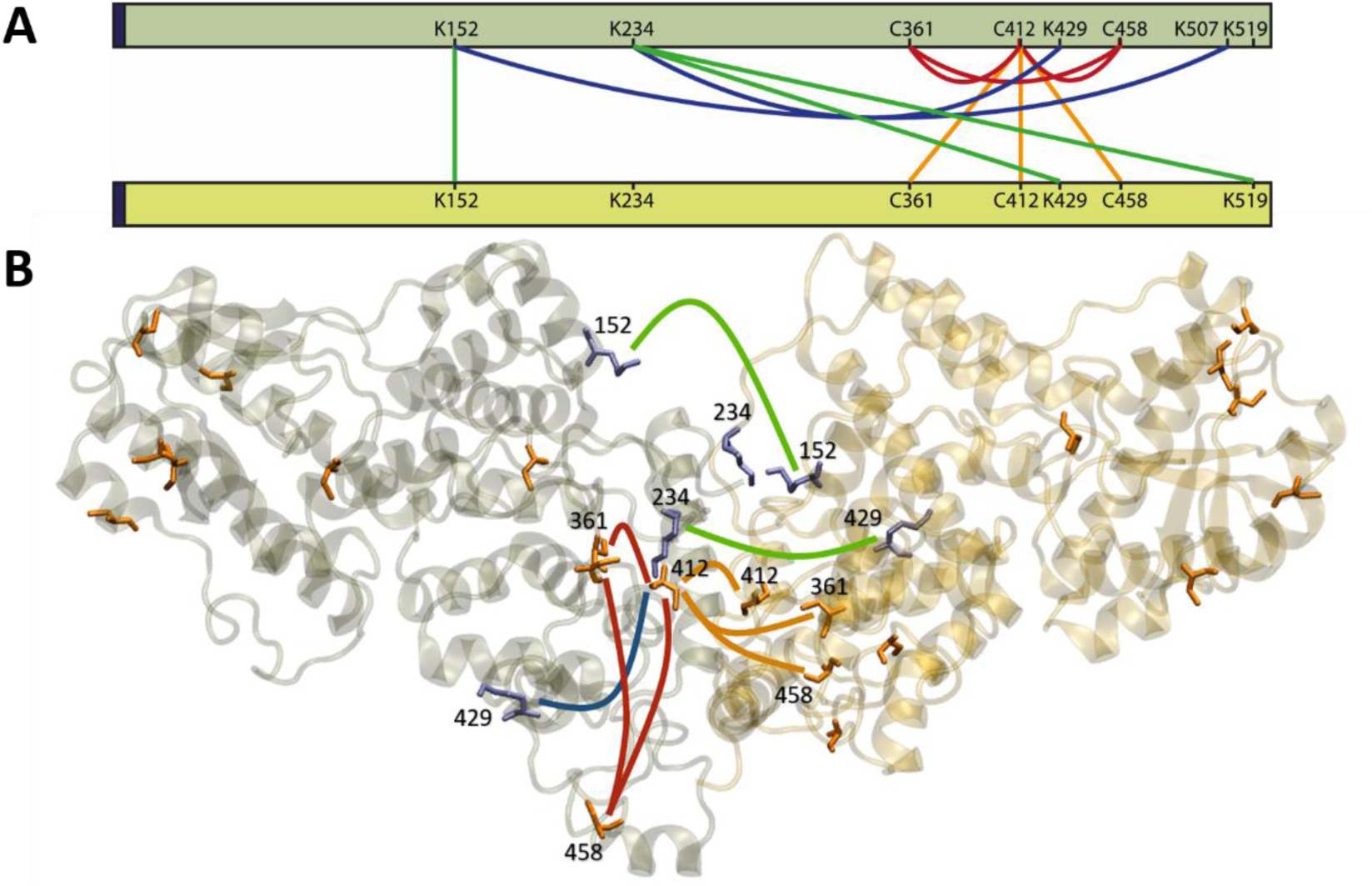
(**A**) Network plot of cross-linking sites found for *Er*Cry4a. The orange and red lines are potential inter-and intramonomer disulphide bonds, respectively. The inter-and intramonomer cross-links formed by DSBU and DSSO are shown in green and blue, respectively. The dark blue region at the N-terminus (left hand side) corresponds to the five additional amino acids that remained after His-tag cleavage. (**B**) Graphical representation of a possible coordination of two *Er*Cry4a monomers. The cross-links are drawn with the same colour code as in (A). Cysteine residues are shown in orange and lysine residues in purple. Only residues 1-495 are displayed.

### Computationally reconstructed *Er*Cry4a dimer structures

As described in the Methods section (Fig. 2), a large number of covalently and non-covalently bound dimers were constructed using ZDOCK and M-ZDOCK. A total of 49 docked structures were generated for a truncated form of *Er*Cry4a (Table 3) using the seven most exposed cysteines as contact residues. Another ten were obtained without specifying which cysteines should be near one another in the docked structure. Twelve of these structures were selected for MD simulations, based on the separations of the cysteine residues (eight with a C412-C412 disulphide, three with C317-C317, one with C189-C485). To these were added: a non-covalently bound structure derived from the crystallographic dimer of mouse Cry2, a dimer with two disulphide bridges, and two structures for the full-length form of *Er*Cry4a, both non-covalent.

Of the 16 dimer structures chosen for MD simulation (see Methods section and Supplementary Material for more details), seven were selected (Table 5) on the basis of their monomer-monomer interaction energies, *E*_tot_: two structures with C412-C412 disulphides (cov^412A^ and cov^412B^), two with C317-C317 bonds (cov^317A^ and cov^317B^), two non-covalent (ncov^A^ and ncov^M^) and the one with two disulphides (cov^D^, C116-C313 and C313-C116). ncov^M^ is the structure based on the crystallographic dimer of *Mm*Cry2.

Various other properties of these structures, derived from the MD simulations, are presented in Table 5 and described below. Of the seven dimers, ncov^A^ and cov^412A^ have the largest internal potential energies (*E*_tot_) and are therefore the most stable. The interaction between the monomers in ncov^A^ is probably stronger because the disulphide bond in cov^412A^ (and the other covalent structures) constrains the monomers at the dimer interface preventing them finding more favourable binding conformations. Note that *E*_tot_ comprises the non-covalent van der Waals and electrostatic interactions between the monomers and does not include the ca. 60 kcal mol^−1^ expected for a disulphide bond^80^. The other dimers featured in Table 5, especially cov^317A^, cov^D^, and cov^317B^, have smaller interaction energies and are therefore less likely to be formed.

**Table 5.** The seven simulated and analysed *Er*Cry4a dimers with the largest non-bonded monomer-monomer interaction energies, *E*_tot._ Results for other dimers are given in the Supplementary Material. The various parameters are discussed in the text.

| Dimer | $-E_{\text{tot}} /$<br>kcal mol<sup>-1</sup> | $R_g / \text{\AA}$ | RMSD/<br>$\text{\AA}$ | RMSF/<br>$\text{\AA}$ | $A_{\text{IS}} / \text{\AA}^2$ | Hydrogen<br>bonds | Salt<br>bridges |
| --- | --- | --- | --- | --- | --- | --- | --- |
| ncov<sup>A</sup> | 857±85 | 35.2±0.3 | 3.5±0.4 | 2.2±1.0 | 2762±292 | 106 | 29 |
| cov<sup>412A</sup> | 526±83 | 41.2±0.2 | 2.5±0.4 | 5.7±1.3 | 1840±115 | 31 | 13 |
| ncov<sup>M</sup> | 505±163 | 33.0±0.3 | 3.4±1.4 | 2.3±1.0 | 2442±284 | 111 | 12 |
| cov<sup>412B</sup> | 437±127 | 41.4±0.7 | 6.2±2.8 | 5.4±1.5 | 1040±310 | 49 | 12 |
| cov<sup>317A</sup> | 186±103 | 38.3±0.3 | 4.0±0.6 | 1.8±0.9 | 1082±220 | 64 | 10 |
| cov<sup>D</sup> | 178±44 | 33.0±0.2 | 2.5±0.3 | 1.3±0.7 | 2140±330 | 47 | 3 |
| cov<sup>317B</sup> | 171±73 | 38.5±0.2 | 3.8±0.8 | 15.6±4.1 | 1070±110 | 59 | 12 |

The stronger binding of the two non-covalent dimers is reflected in their smaller radius of gyration, *R*_g_, which is a measure of the compactness of the structure. The cov^D^ dimer also has a relatively small *R*_g_ presumably because of the constraining effects of its two disulphide bonds.

Another indication of protein stability is provided by the average root-mean-squared deviation (RMSD) of the backbone atom positions from those in a reference structure immediately prior to the production simulation^81^. Smaller values of this parameter correspond to dimers in which the initial docked structure did not alter significantly throughout the simulation. Apart from cov^412B^ (6.2 Å), all the dimers in Table 5 have RMSDs less than 4 Å. Dimers cov^412A^ and cov^D^ (RMSD ≍ 2.5 Å) are the most stable as judged by this measure.

A parameter that quantifies the degree of internal mobility of the dimer is the root-mean-square amplitude of the structural fluctuations (RMSF) during the course of an MD simulation. RMSF values were calculated by comparing the positions of each carbon atom at every simulation step to its position in the average structure. With the exception of cov^317B^, which is much less rigid than the others, all the structures in Table 5 have comparable values of this parameter. Certain regions of the monomers are more flexible than others, in particular the C-terminal extension (residues 470-495) and the phosphate-binding loop (residues 231-248). For the most part, this flexibility seems to be retained on dimerisation.

A further measure of the interaction between the monomer units of a dimer is the interaction surface area, *A*_IS_ (Table 5), defined as the difference between the solvent accessible surface areas of the separate monomers and that of the dimer. Larger values of *A*_IS_ should correlate with stronger binding energies (-*E*_tot_) as seems to be broadly the case for the dimers in Table 5. Two additional parameters are the number of hydrogen bonds and the number of salt bridges that link the two components of the dimer. Only salt bridges present in more than 10% of the MD frames were counted. All values have been averaged over the duration of the corresponding MD simulations. The two non-covalent structures have the largest interaction surface areas and many more hydrogen bonds than the covalently bound dimers.

Figures 7A and B show the dimer interfaces of cov^412A^ and ncov^A^. Consistent with the data in Table 5, the non-covalent dimer has a more extensive interface.

**Figure 7.**
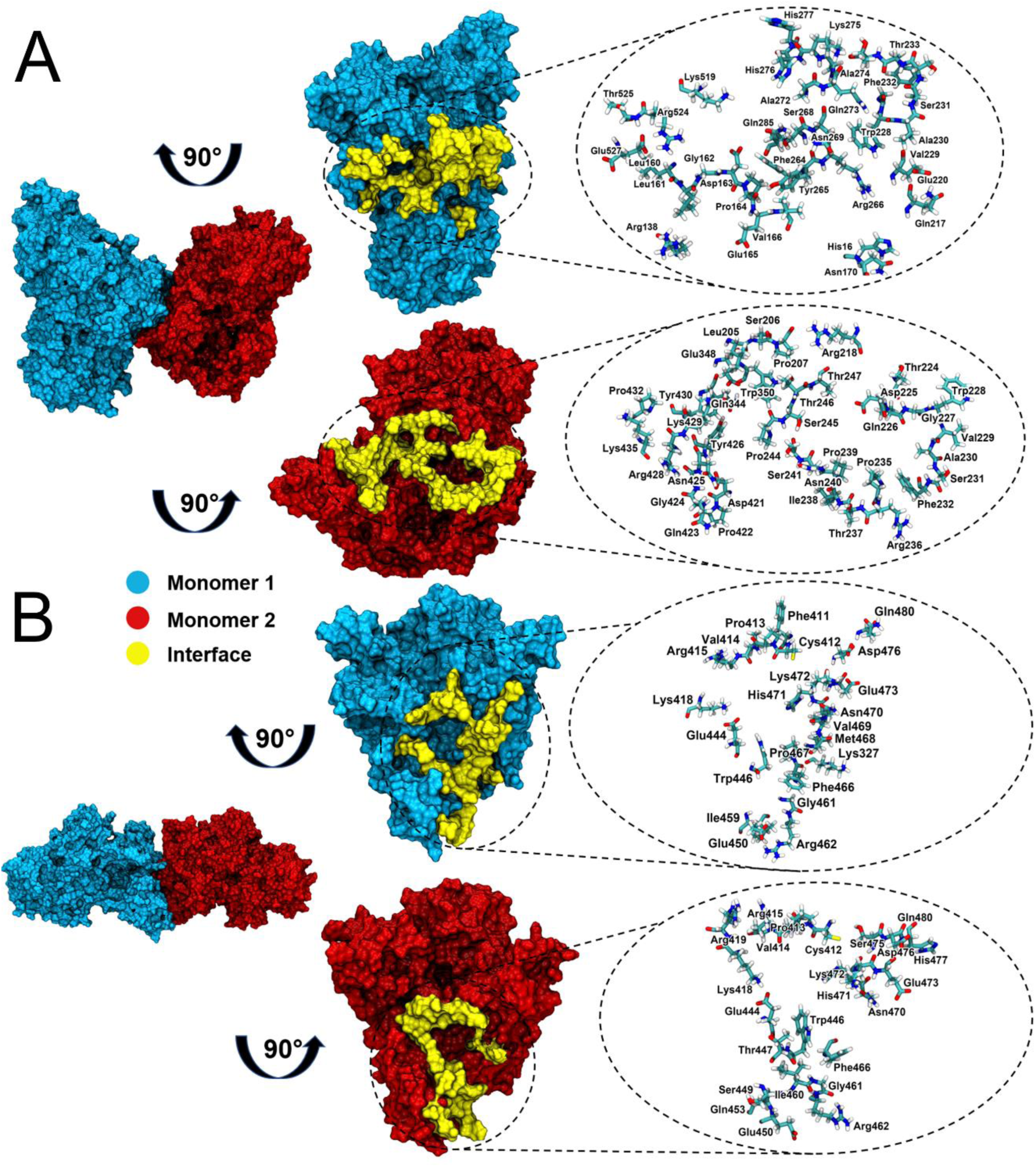
The dimer interfaces of (**A**) ncov^A^ and (**B**) cov^412A^. Left: dimer structures. Middle: 90°-rotated monomers showing the interacting residues (yellow). Right: representations of the contact residues in the interface.

As described in the Methods section, none of the docked dimers obtained using C68, C73, C116 and C179 as the contact residue were deemed worth simulating by MD. These four residues, together the four with the smallest solvent accessibilities (C257, C313, C361, C458) appear to have a low propensity to dimerise via the formation of a single disulphide bond. All of the C189-linked dimers which satisfied our conditions for MD simulation had smaller binding energies than the seven in Table 5 and therefore appear less likely to be formed. Of the covalently bound dimers, this leaves C317-C317, C412-C412 and the double-disulphide dimer, cov^D^, which involves C116 and C313. Based on the data in Table 5, C412 seems most likely to form covalent links between *Er*Cry4a monomers. The other main conclusion that can be drawn from these studies is that the possibility of non-covalently bound dimers cannot be dismissed.

## Conclusions

The experiments described here show that *Er*Cry4a readily forms dimers and that they are covalently linked by disulphide bonds. Several cysteines seem to be involved, C317 and C412 being the most likely. Computational modelling supports this conclusion, provides insights into the relative orientation of the monomer units, and suggests that non-covalently bound dimers may also be relatively stable. The relevance of these findings to the proposed role of Cry4a in avian magnetoreception is unclear.

That *Er*Cry4a dimers are not readily disrupted in a reducing environment (10 mM DTT, Fig. 3) suggests that they could persist *in vivo*. Despite the reducing environment of the cell, there are well-documented examples of biologically significant intra-cellular disulphide-linked proteins involved in redox processes^82–84^. Moreover, little is known about the redox conditions in the avian photoreceptor cells that are thought to contain the magnetoreceptors. It therefore seems premature to exclude the possibility that *Er*Cry4a dimerisation could have a biological function.

If *Er*Cry4a does have a role in magnetic sensing and/or signalling, then an immediate question is why the protein has evolved to stabilise inter-monomer disulphide bonds to the extent that only a proportion of the proteins dimerise under the conditions of the experiments reported here. One explanation is that these dimers are simply aggregates of unfolded monomers. However, we think this is unlikely, as the *Er*Cry4a dimers bind FAD and have native MS charge-state distributions that are similar to those of the monomers suggesting that the protein remains correctly folded and therefore potentially functional in the dimeric state. Partially unfolded proteins typically display higher charge states than the native states. An alternative explanation is that a monomer-dimer equilibrium could play a regulatory role in magnetic sensing or signal transduction. Activation of plant cryptochromes 1 and 2 leads to homo-oligomerisation^3, 5, 40, 41, 85^. Additionally, the formation of a disulphide bond influences the interaction between mouse cryptochrome 1 and the circadian protein Period (PER) and depends on the metabolic/oxidative state inside the cell^86^. Taken together, these examples show that photoactivation of some cryptochromes causes oligomerisation and that cryptochrome’s affinity for other proteins could rely on disulphide bond formation.

An observation reported by Xu et al.^15^ (in their Fig. S5) may be relevant here. Samples of *Er*Cry4a that had been purified without adding 10 mM BME to prevent dimerisation showed clear evidence for the formation of relatively long-lived (nanosecond to microsecond) photo-excited singlet (^S^FAD*) and triplet (^T^FAD*) states of FAD in addition to the normal photo-induced electron transfer reactions that generate radical pairs. It was speculated that subtle differences in protein conformation, induced by dimerisation, might inhibit electron transfer from the tryptophan tetrad to ^S^FAD*, thereby stabilising ^S^FAD* and allowing formation of ^T^FAD* by intersystem crossing. Samples of *Er*Cry4a purified with 10 mM BME, and therefore more likely to be monomeric, showed much weaker signals from ^S^FAD* and ^T^FAD*.

While recognising that the experiments described here have not been performed in cells, they do show it is quite possible that dimerisation could have an impact on the *in vivo* function of avian cryptochromes and provide an incentive for further work in this area.

## Supporting information

Supplementary figures S1-S14 and Tables S1-S20 generated in the present study.

## Supporting Information

Supplementary figures S1-S14 and Tables S1-S20 generated in the present study. Additional supplementary data and protocols used in the present investigations are provided.

## Acknowledgments

The authors would like to thank the Volkswagen Foundation (Lichtenberg Professorship to IAS), the Deutsche Forschungsgemeinschaft (DFG; GRK1885: Molecular Basis of Sensory Biology and SFB 1372: Magnetoreception and Navigation in Vertebrates), the Ministry for Science and Culture of Lower Saxony (Simulations Meet Experiments on the Nanoscale: Opening up the Quantum World to Artificial Intelligence (SMART) and Dynamik auf der Nanoskala: Von kohärenten Elementarprozessen zur Funktionalität (DyNano)), the Japan Science and Technology Agency (JST) PRESTO (Quantum Bio) Grant No. JPMJPR19G1 (LMA), the European Research Council (under the European Union’s Horizon 2020 research and innovation programme, Grant Agreement No. 810002, Synergy Grant: *QuantumBirds*), and the Office of Naval Research Global, award no. N62909-19-1-2045. Computational resources for the simulations were provided by the CARL Cluster at the Carl-von-Ossietzky University, Oldenburg, supported by the DFG and the Ministry for Science and Culture of Lower Saxony. The authors gratefully acknowledge the computing time granted by the Resource Allocation Board and provided on the supercomputer Lise and Emmy at NHR@ZIB and NHR@Göttingen as part of the NHR infrastructure. The calculations for this research were conducted with computing resources under the project nip00058. TJE was supported by funds from the Royal Society under a Royal Society Newton International Fellowship. ASG is grateful to Mark Pepys who provided CRP for the mass spectrometric investigation and to Josh Bishop who helped with the mass photometry measurements and data analysis.

