## Supplementary figures S1-S14 and Tables S1-S20 generated in the present study. for "Dimerisation of European robin cryptochrome 4a"

### Dimerisation of European robin cryptochrome 4a (Supplementary Material)

### Table of Contents


Experimental supplementary figures ........ S3

Full SDS-PAGE gel, peak isolation using tandem native MS and XL-MS results ........ S3

Photometric cysteine exposure measurements ........ S4

Experimental supplementary tables ........ S5

Computational supplementary material ........ S7

*Er*Cry4a non-covalent dimers ........ S8

*Er*Cry4a 317 dimer family ........ S11

*Er*Cry4a cov$^{D}$ – a dimer with two disulphide bonds ........ S13

*Er*Cry4a 189 dimer family ........ S15

*Er*Cry4a 412 dimer family ........ S17

Distance analysis between cysteine residues ........ S22

Molecular dynamics simulation protocol ........ S31

References ........ S32

### Experimental supplementary figures

#### Full SDS-PAGE gel, peak isolation using tandem native MS and XL-MS results

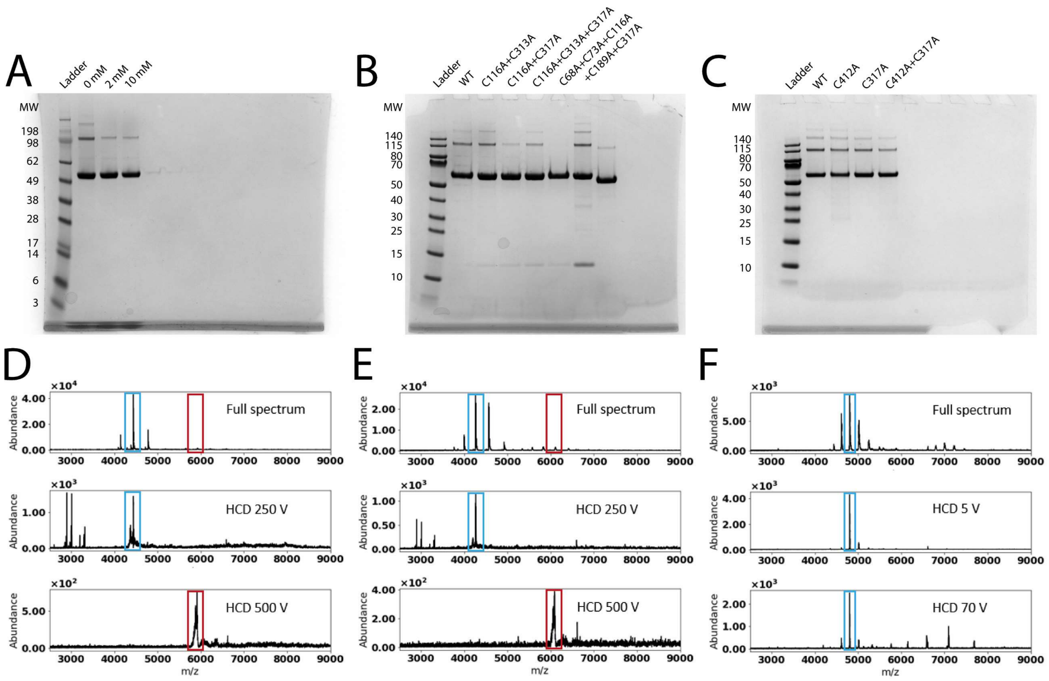


**Figure S1**: (**A**)-(**C**) Denaturing SDS-PAGE gels run for the different *Er*Cry4a proteins displayed in the native MS spectra shown in the main text. Proteins are as indicated at the tops of the columns. All leftmost columns display protein ladders. In (**A**) the SeeBlue™ Plus2 Pre-stained Protein Standard ladder (Invitrogen) was used and in (**B**) and (**C**) the PageRuler Prestained Protein Ladder (Thermo Scientific) was used with the molecular weight (MW) mass markers in kDa as indicated. The columns displayed with no indicator are not relevant for the present investigation. (**D**)-(**F**) Native MS in combination with tandem MS was used to isolate a monomer and a dimer peak of *Er*Cry4a WT (**D**) (without its His-tag) and *Er*Cry4a C317S (**E**) (with His-tag; shipped in 10 mM BME to prevent higher order oligomerisation during transport) to compare their stability upon exposure to high HCD (high energy collisional dissociation) energies. The first rows of both (**D**) and (**E**) show the full spectra, the second the isolated dimer peaks and the third the isolated monomer peaks. HCD values were applied as indicated in the spectra. The spectra displayed for comparison in (**F**) were of CRP (C-reactive protein), a pentameric protein known to be non-covalently bound[1]. The first row shows the full spectrum, the second an isolated pentamer peak and the third shows the isolated pentamer peak after applying an HCD value of 70 V, displaying how it falls apart into smaller subunits at comparatively low HCD energies. The *Er*Cry4a dimers still did not dissociate when the highest HCD energies possible on the instrumental setup (500 V) were applied. The coloured rectangles show which peaks were isolated using tandem MS.

#### Photometric cysteine exposure measurements

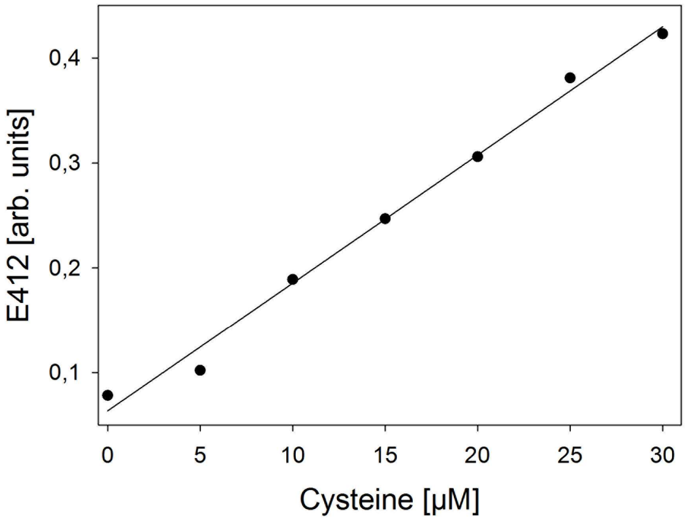


**Figure S2**: Cysteine accessibility calibration curve. 60 µM of DTNB were added to 0, 5, 10, 15, 20, 25 and 30 µM of L-cysteine. After 10 min incubation at room temperature the absorbance was measured at 412 nm. The *y*-axis shows the absorbance of TNB at 412 nm resulting from the reaction of DTNB with the thiol groups of the cysteines. The solid line comes from a linear regression analysis and has slope 0.0124 $\mu M^{-1}$.

### Experimental supplementary tables

**Table S1**: Absorbance of 5 µM *Er*Cry4a wildtype (WT) and its mutants measured at 412 nm after 10 min incubation with 60 µM DTNB and the corresponding number of accessible cysteines.

| WT absorption | Accessible cysteines | C317A absorption | Accessible cysteines | C412A absorption | Accessible cysteines | C116-C317 absorption | Accessible cysteines |
|---|---|---|---|---|---|---|---|
| 0.30 | 4.77 | 0.26 | 4.13 | 0.27 | 4.17 | 0.16 | 3.12 |
| 0.30 | 4.76 | 0.26 | 4.09 | 0.27 | 4.14 | 0.17 | 3.30 |
| 0.28 | 4.58 | 0.28 | 4.34 | 0.27 | 4.14 | 0.18 | 3.35 |
| 0.30 | 4.82 | 0.28 | 4.30 | 0.27 | 4.25 | 0.17 | 3.30 |
| 0.31 | 4.98 | 0.28 | 4.31 | 0.26 | 4.09 | 0.18 | 3.43 |

**Table S2**: XL-MS results. Only disulphide bonds and crosslinks between peptides containing lysine residues[2] that were identified with a MeroX score greater than 50 were considered candidates for further analysis. The residues in brackets clarify the position of the linked residue within each peptide.

| Link | Experiment | Score | Peptide 1 | Peptide 2 | Linked residues |
|---|---|---|---|---|---|
| DSBU | 1 (dimer) | 81 | 144-158 (K14) | 144-158 (K14) | K152-K152 |
| | | 76 | 224-241 (K16) | 434-440 (K1) | K234-K429 |
| DSBU | 1 (monomer) | 59 | 224-241 (K16) | 434-440 (K1) | K234-K429 |
| DSSO | 1 (dimer) | 52 | 224-241 (K16) | 516-526 (K9) | K234-K519 |
| DSSO | 1 (monomer) | 109 | 144-158 (Y13) | 503-514 (K10) | K152-K507 |
| disulphide | 1 (dimer) | 144, 106, 90 | 415-420 (C3) | 415-420 (C3) | C412-C412* |
| | | 103, 97 | 362-370 (C5) | 415-420 (C3) | C361-C412 |
| | | 120 | 460-467 (C4) | 415-420 (C3) | C458-C412 |
| disulphide | 1 (monomer) | 121, 97, 90, 79 | 415-420 (C3) | 415-420 (C3) | C412-C412* |
| | | 117 | 460-467 (C4) | 415-420 (C3) | C458-C412 |
| disulphide | 2 (monomer) | 145, 101, 97, 94, 93 | 362-370 (C5) | 415-420 (C3) | C361-C412 |
| | | 164, 149, 81 | 415-420 (C3) | 415-420 (C3) | C412-C412 |
| | | 166, 109 | 460-467 (C4) | 415-420 (C3) | C458-C412 |
| | | 95 | 362-370 (C5) | 460-467 (C4) | C361-C458 |

*A disulphide bond between two C412 residues was found in both the monomer and dimer fractions, possibly due to cross-contamination between the two fractions on the SDS-PAGE gel.

### Computational supplementary material

**Table S3**. Solvent exposure and locations of the cysteine residues in WT *Er*Cry4a calculated as described in the Methods section and Eq. (1) in the main text. The values were averaged over the duration of the production MD simulation.

| Cysteine | Secondary structure | Solvent exposure / % |
|---|---|---|
| 317 | unspecified turn | 71.9 |
| 116 | α-helix | 57.5 |
| 189 | α-helix | 55.5 |
| 68 | α-helix | 44.7 |
| 412 | coil, close to α-helix | 30.6 |
| 73 | coil, close to β-strand | 22.5 |
| 179 | coil, close to α-helix | 6.5 |
| 313 | unspecified turn | 6.1 |
| 458 | coil | 1.7 |
| 361 | α-helix | 1.1 |
| 257 | unspecified turn | 0.2 |

#### *Er*Cry4a non-covalent dimers

The full length *Er*Cry4a WT structure was used to produce stable non-covalently bound dimers (illustrated in Fig. S3). Several analyses were performed on the simulated dimers: the results are summarised in Table S4 and Fig. S4. The average $\overline{RMSD}$ values calculated over the duration of the production simulation are shown in Table S4, while the time evolution of the $\overline{RMSD}$ is presented in Fig. S4. The $ncov^{A}$ and $ncov^{M}$ dimers appear to be rather stable with an average $\overline{RMSD}$ value of 3.51 ± 0.36 Å and 3.36 ± 1.36 Å, respectively, whereas $ncov^{4}$ is rather unstable with an average $\overline{RMSD}$ value of 8.98 ± 2.00 Å. The contrast in the stabilities of the three dimers is attributed to the differences in interaction energy and the hydrogen bonding network, both factors being much stronger in $ncov^{A}$ and $ncov^{M}$ (see Table S4).

Average interaction energies, $E_{tot}$, for the non-covalent dimer family are given in Table S4. The $ncov^{A}$ dimer was selected for further comparative analysis because of its exceptionally strong interaction energy that already manifests itself after the 2 ns equilibration simulation ($E_{tot}$ = −928 ± 25 kcal mol$^{-1}$, Table S5). The $ncov^{4}$ dimer was considered interesting because of its similar spatial arrangement to the covalent dimer $cov^{317A}$ discussed below. Considering that the full length *Er*Cry4a protein was used for $ncov^{4}$ and $ncov^{A}$ simulations, the presence of the CTT might have added to the stability of the dimer by contributing favourably to the resulting interaction energies.

The average value of the radius of gyration, $R_g$, was computed for the non-covalent dimers, where the averaging was over the span of the MD trajectories. The results in Table S4 reveal that $R_g$ for $ncov^{A}$ and $ncov^{M}$ is significantly lower than for $ncov^{4}$, which indicates that the protein structures are more compact.

The average $\overline{RMSF}$ value turns out to be smallest for $ncov^{A}$ (Table S4), where the largest fluctuations occur in the CTT domain (residues 498-527) as one would expect for an intrinsically disordered region of the protein (see Fig. S4D-F).

There are 106 and 111 inter-monomer hydrogen bonds in the $ncov^{A}$ and $ncov^{M}$ dimers, respectively, while only 40 exist in $ncov^{4}$. The numbers of inter-monomer salt bridges in $ncov^{4}$ and $ncov^{A}$ are similar and significantly larger than in $ncov^{M}$.

The interaction surface areas of $ncov^{A}$ and $ncov^{M}$ are more than double that of $ncov^{4}$. Interestingly, even without the CTT, $ncov^{M}$ has a bigger interaction surface than $ncov^{4}$. Taking all the factors into consideration, $ncov^{A}$ and $ncov^{M}$ were selected as the two most promising non-covalent *Er*Cry4a candidates for a further comparative analysis.

**Table S4**: Summary of the characteristics of the non-covalent dimers. The table includes the computed values of the radius of gyration $R_g$, the average $\overline{RMSD}$ and $\overline{RMSF}$ values, the total area of the binding interface $A_{IS}$, and the total number of inter-monomer hydrogen bonds and salt bridges. Only salt bridges present in more than 10% of the MD frames were counted. All values have been averaged over the duration of the corresponding MD simulations.

| Dimer | $-E_{tot}$ / kcal mol$^{-1}$ | $R_g$ / Å | $\overline{RMSD}$/Å | $\overline{RMSF}$/Å | $A_{IS}$ / Å$^2$ | Hydrogen bonds | Salt bridges |
|---|---|---|---|---|---|---|---|
| ncov$^4$ | 426±95 | 41.8±0.9 | 9.0±2.0 | 3.2±1.5 | 1125±161 | 40 | 33 |
| ncov$^A$ | 857±85 | 35.2±0.3 | 3.5±0.4 | 2.2±1.0 | 2762±292 | 106 | 29 |
| ncov$^M$ | 505±163 | 33.0±0.3 | 3.4±1.4 | 2.3±1.0 | 2442±284 | 111 | 12 |

**Table S5**: Ten most favourable non-covalently bound full length *Er*Cry4a dimers, ncov$^n$, produced by ZDOCK. Average interaction energies $E_{tot}$ between each monomeric subunit after 2 ns equilibration are shown together with the van der Waals, $E_{vdw}$, and Coulomb, $E_{elec}$, contributions.

| Dimer | $-E_{elec}$ / kcal mol$^{-1}$ | $-E_{vdW}$ / kcal mol$^{-1}$ | $-E_{tot}$ / kcal mol$^{-1}$ |
|---|---|---|---|
| ncov$^1$ | 320±61 | 91±8 | 411±65 |
| ncov$^2$ | 237±47 | 114±12 | 351±42 |
| ncov$^3$ | 332±41 | 118±13 | 450±36 |
| ncov$^4$ | 245±37 | 73±13 | 318±33 |
| ncov$^5$ | 106±44 | 65±8 | 171±41 |
| ncov$^6$ | 513±41 | 111±10 | 624±42 |
| ncov$^7$ | 245±42 | 102±8 | 347±43 |
| ncov$^8$ | 283±29 | 55±7 | 338±27 |
| ncov$^9$ = ncov$^A$ | 826±66 | 102±8 | 928±25 |
| ncov$^{10}$ | 319±24 | 51±4 | 370±25 |

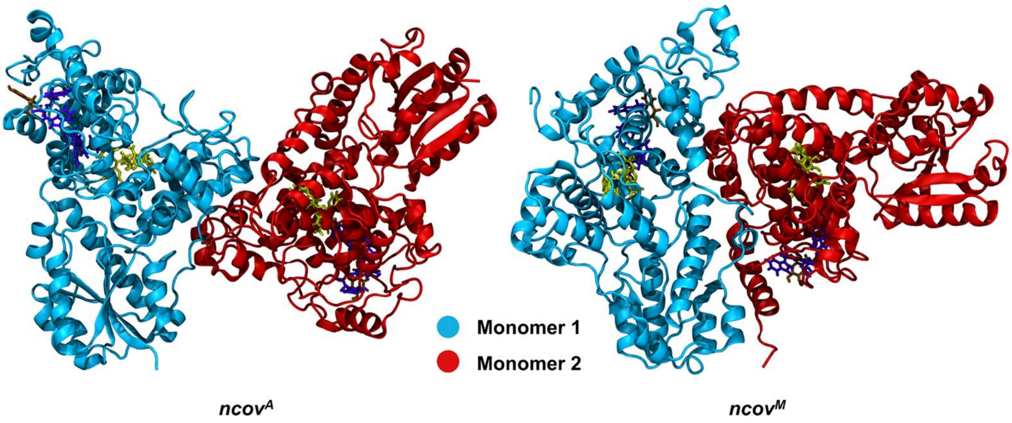


**Figure S3**: Structures of the non-covalently bound dimers ncov$^{A}$ and ncov$^{M}$. The FAD cofactor, Trp-tetrad (W395, W372, W318, W369), and Y319 are shown in yellow, violet and ochre, respectively.

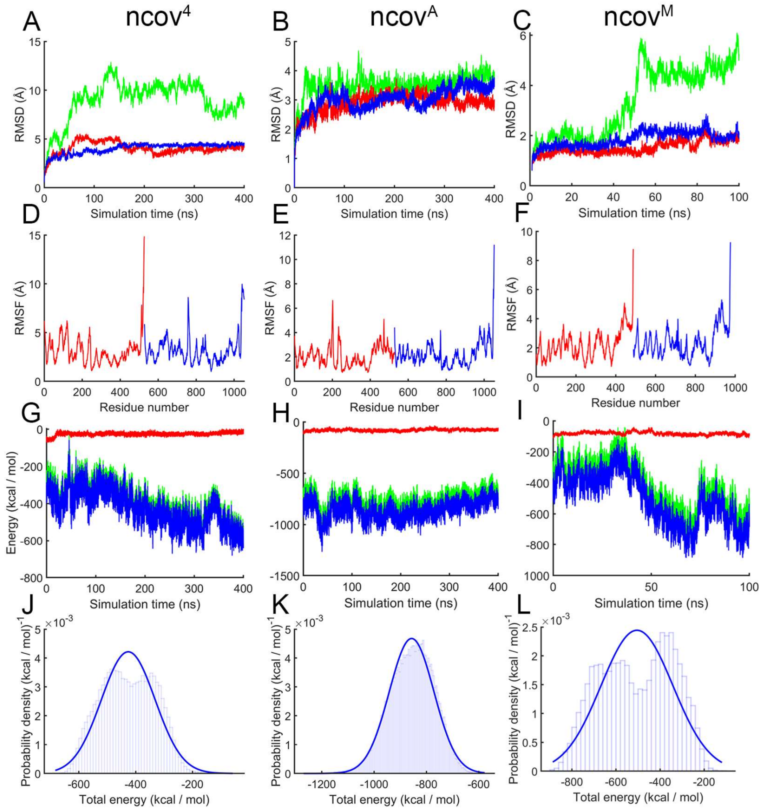


**Figure S4**: Time-evolution of the root mean square displacement (RMSD) values computed for the backbone atoms of the non-covalently bound dimers ncov$^{4}$, ncov$^{A}$ and ncov$^{M}$ during the 400 ns production simulation (**A, B, C**). Monomers (blue and red); dimer (green line). Average root mean square fluctuations $\overline{RMSF}$ computed for the residues of the non-covalently bound dimers ncov$^{4}$, ncov$^{A}$ and ncov$^{M}$ (**D, E, F**). Red corresponds to monomer A and blue to monomer B. (**G, H, I**) Time-evolution of the interaction energies for non-covalently bound dimers ncov$^{4}$, ncov$^{A}$ and ncov$^{M}$. Red and green denote respectively the van der Waals and Coulomb contributions to the total interaction energy between the monomers, shown in blue. (**J, K, L**) Probability density distributions of the interaction energies.

#### *Er*Cry4a 317 dimer family

Residue C317 is of interest in the context of covalent dimerisation of *Er*Cry4a as it is close to the Trp tetrad (which is involved in magnetic sensing) and is the most solvent-exposed cysteine residue in WT *Er*Cry4a (Fig. 4A and Table S3). C317 is also considered a promising linking residue, as a result of the experiments described in the main text.

Three different dimeric structures were constructed to investigate the involvement of C317 in covalently-bound *Er*Cry4a dimers (see Table S16). Dimers $cov^{317A}$ and $cov^{317B}$ are illustrated in Fig. S5. Table S6 gives the average $\overline{RMSD}$ values for the three dimers covalently linked through the C317 residue. The time-dependence of the RMSD is given in Fig. S6A-C. The results in Table S6 demonstrate that $cov^{317A}$ and $cov^{317B}$ are much more stable than $cov(317)^3$. Table S6 also gives the average $\overline{RMSF}$ values which provide information on the flexibility of the three structures. The average $\overline{RMSF}$ is smallest for $cov^{317A}$, indicating that the structure is least flexible, and unusually large for $cov^{317B}$.

The analysis of the average radius of gyration, $R_g$, in Table S6 shows that $R_g$ is comparable for all three structures indicating that their geometric shapes are similar, which is also suggested by their similar interaction surface areas. Furthermore, the interaction energy of the subunits shown in Fig. S6D-F, shows that the most favourable interactions occur between the subunits of the $cov^{317A}$ dimer. The hydrogen bonding network is more elaborate for the $cov(317)^3$ dimer; here the number of hydrogen bonds could be directly related to the larger interaction surface area. The salt bridge analysis favours $cov^{317A}$, supporting this dimer as the representative candidate of the *Er*Cry4a C317 dimer family.

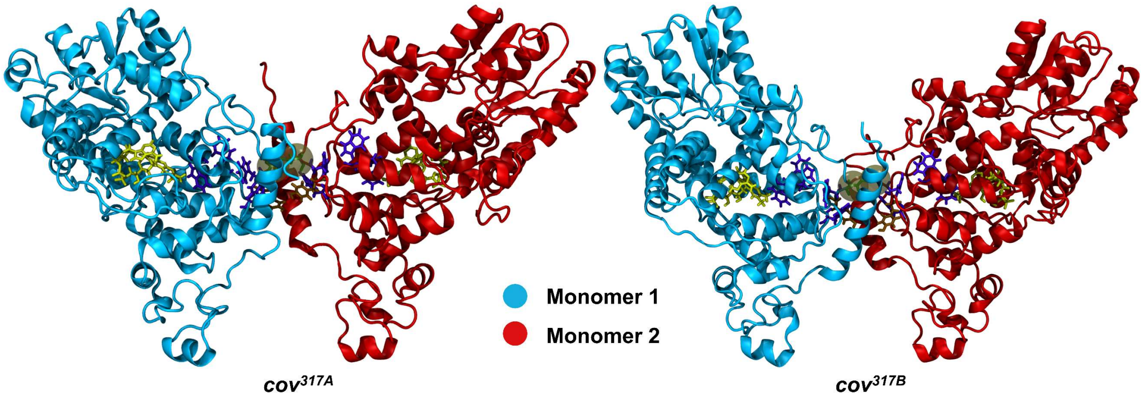


**Figure S5**: Structures of the covalently bound dimers of *Er*Cry4a, $cov^{317A}$ and $cov^{317B}$. The FAD cofactor, Trp-tetrad (W395, W372, W318, W369), and Y319 are shown in yellow, violet and ochre, respectively. Brown spheres indicate the approximate position of the C317-C317 disulphide bond between the monomers.

**Table S6**: Summary of the characteristics of the cov(317)$^n$ dimer family. The table includes the computed values of the radius of gyration $R_g$, the average $\overline{RMSD}$ and $\overline{RMSF}$ values, the total area of the binding interface $A_{IS}$, as well as the total number of inter-monomer hydrogen bonds and salt bridges. Only salt bridges present in more than 10% of the MD frames were counted. All values have been averaged over the duration of the corresponding MD simulations.

| Dimer | $-E_{tot}$ / kcal mol$^{-1}$ | $R_g$ / Å | $\overline{RMSD}$ / Å | $\overline{RMSF}$ / Å | $A_{IS}$ / Å$^2$ | Hydrogen bonds | Salt bridges |
|---|---|---|---|---|---|---|---|
| cov$^{317A}$ | 186±103 | 38.3±0.3 | 4.0±0.6 | 1.8±0.9 | 1082±220 | 64 | 10 |
| cov$^{317B}$ | 171±73 | 38.5±0.2 | 3.8±0.8 | 15.6±4.1 | 1070±110 | 59 | 12 |
| cov(317)$^3$ | −42±96 | 35.7±1.3 | 8.2±2.9 | 10.9±2.9 | 1173±198 | 83 | 6 |

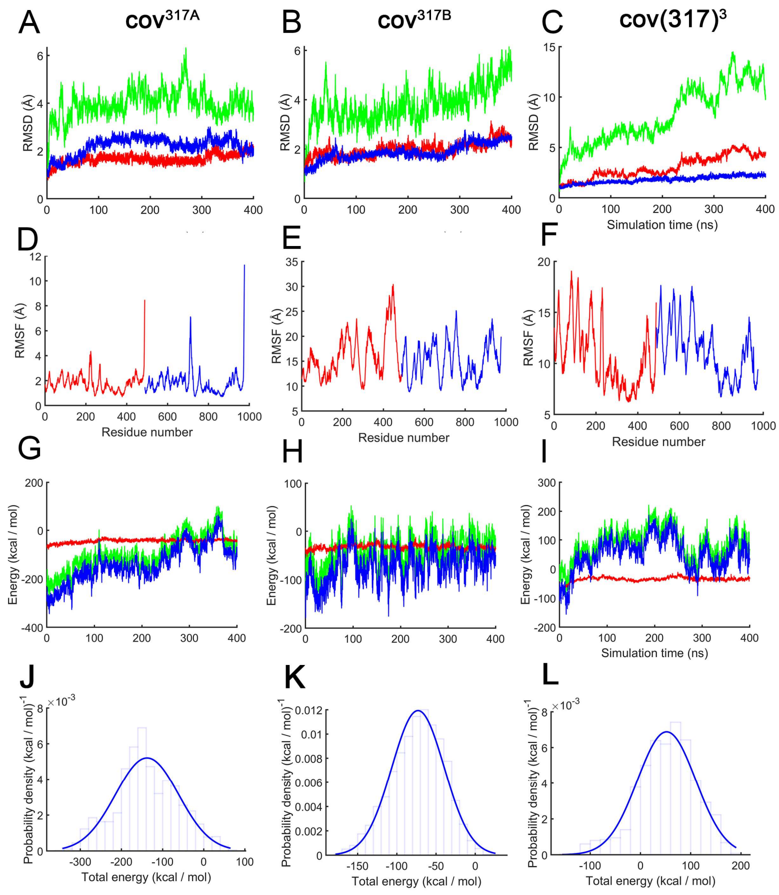


**Figure S6**: Time evolution plots for the *Er*Cry4a C317 dimer family. (**A-C**): Root mean square deviation (RMSD) of the backbone atoms of three members of the *Er*Cry4a C317 dimer family, computed for the dimers (green) and monomers (red and blue), after equilibration. (**D-F**): RMSF values shown per residue. Residues were represented by C$_\alpha$ atoms. (**G-I**): Time evolution of the interaction energies between two monomeric subunits of the *Er*Cry4a dimers. Red and green denote respectively the van der Waals and Coulomb contributions to the total interaction energy between the monomers, shown in blue. (**J-L**): Probability density distributions of the interaction energies.

#### *Er*Cry4a cov$^D$ – a dimer with two disulphide bonds

Figure S1 shows that *Er*Cry4a dimers can still be formed after mutation of important cysteine residues that account for a part of the dimerisation. Fig. S1 also indicates that small amounts of higher order oligomers exist in addition to the more prominent monomers and dimers, suggesting that several oligomerisation surfaces in *Er*Cry4a may exist and therefore that more than one disulphide bridge could be involved in linking monomers. Oligomers were also found for *Arabidopsis thaliana* cryptochrome 2 (*At*Cry2) where a tetrameric structure is important in the regulation of plant growth[3,4]. Another experimental indication of dimer stability involves the use of higher HCD which demonstrates that *Er*Cry4a dimers do not disintegrate easily (Fig. S1). This result suggests that a covalent linkage between the subunits might involve more than just one disulphide bond. Using M-ZDOCK[5], a tool that symmetrically docks multimers, a dimeric structure was found in which C116 in each of the monomers was close to C313 in the other monomer (see Table S19). This led the construction of a potential stable *Er*Cry4a dimer containing two covalent bonds between the monomers, cov$^D$ = cov(116$^A$313$^B$ - 313$^A$116$^B$), where A and B stand for monomers (Fig. S7).

Table S7 shows the $\overline{RMSD}$ values indicating that the dimeric cov$^D$ structure is stable. A full temporal analysis of *RMSD* and *RMSF* is shown in Fig. S8. Furthermore, the $\overline{RMSF}$ analysis indicates low flexibility, while the interaction energy between the two subunits, averages at $-178 \pm 44$ kcal mol$^{-1}$ and is comparable with the values for cov$^{317A}$ and cov$^{317B}$, even though the interaction surface area for cov$^D$ is twice those of the cov(317) dimers. Figure S7 shows that the *Er*Cry4a cov$^D$ dimer has an inversion centre (centre of symmetry) which is not found for any of the dimeric structures presented in Fig. 1 in the main text.

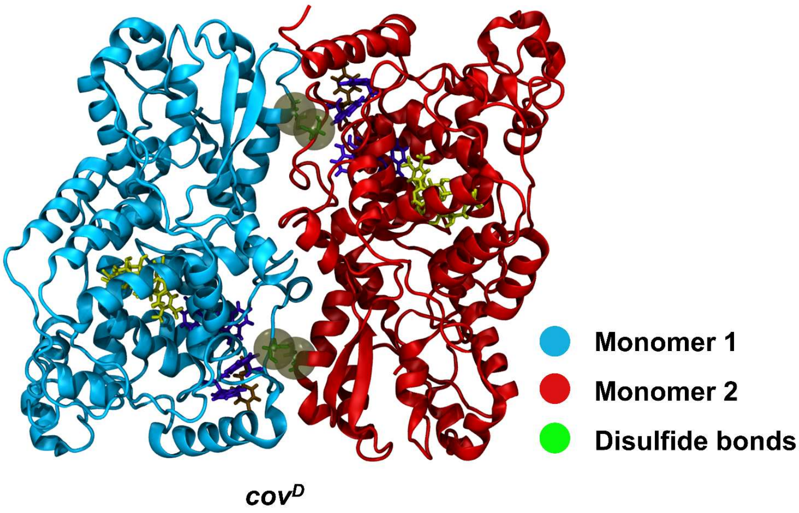


**Figure S7**: Structure of the doubly covalently bound *Er*Cry4a cov$^D$ dimer. The FAD cofactor, Trp-tetrad (W395, W372, W318, W369), and Y319 are shown in yellow, violet and ochre, respectively. Brown spheres indicate the approximate position of the C116-C313 disulphide bonds between the monomers.

**Table S7**: Summary of the analysis of the cov$^D$ dimer similar to the results shown in Tables S4 and S6. Interaction energy values shown here account only for non-bonded interaction between the two monomeric subunits. Only those hydrogen bonds appearing between two subunits are counted. $A_{IS}$ is the interaction surface area between the two monomeric subunits.

| Dimer | $-E_{tot}$ / kcal mol$^{-1}$ | $R_g$ / Å | $\overline{RMSD}$ / Å | $\overline{RMSF}$ / Å | $A_{IS}$ / Å$^2$ | Hydrogen bonds | Salt bridges |
|---|---|---|---|---|---|---|---|
| cov$^D$ | 178 ± 44 | 33.0 ± 0.2 | 2.5 ± 0.3 | 1.3 ± 0.7 | 2140 ± 330 | 47 | 3 |

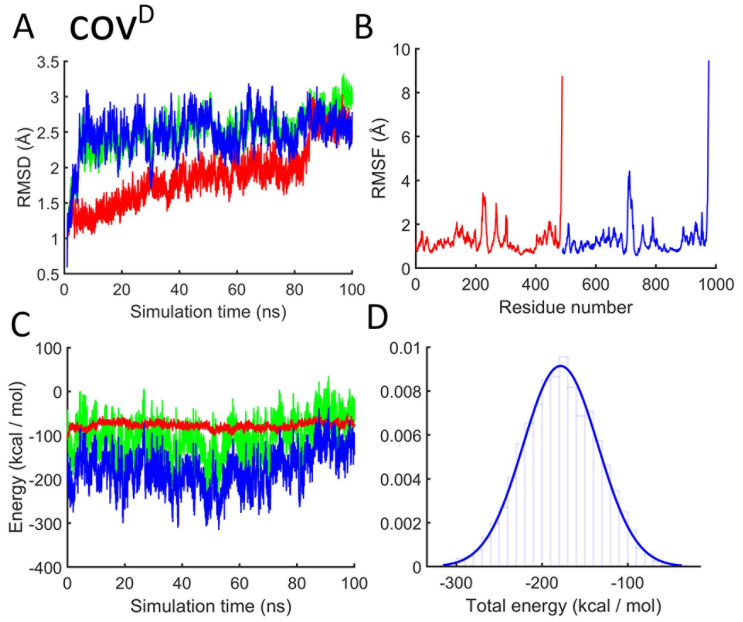


**Figure S8**: Time evolution of the root mean square deviation (RMSD) of the backbone atoms of the *Er*Cry4a cov$^D$ dimer, computed relative to the structure of the corresponding dimer after equilibration (**A**). Average root mean square fluctuations $\overline{RMSF}$ values computed for the residues of the cov$^D$ dimer (**B**). Red corresponds to monomer A and blue to monomer B. (**C-D**): Time evolution of the interaction energies between two monomeric subunits of the cov$^D$ *Er*Cry4a dimer and the corresponding probability density distribution of the interaction energy. Red and green denote respectively the van der Waals and Coulomb contributions to the total interaction energy between the monomers, shown in blue.

#### *Er*Cry4a 189 dimer family

Cys189, the third most solvent-exposed residue of monomeric *Er*Cry4a, was found to have a possible binding motif to the Cys458 residue from another monomer, suggesting a dimer with a disulphide bond between the Cys189 and Cys458, denoted $cov^{189A}$ (see Table S15).

Computational analysis reveals that $cov^{189A}$ is stable, with a low RMSF. Although its interaction surface is not particularly large compared to some of the other covalent dimers, the interaction energy of its monomers, −533 kcal mol$^{-1}$, is the largest of all the covalent dimers studied. Figure S9 shows the spatial orientation of the monomeric subunits; this structure does not have inversion symmetry. Figure S10 shows the temporal analysis of the RMSD, RMSF and $E_{tot}$.

**Table S8**: Summary of the analysis of the $cov^{189A}$ dimer. $E_{tot}$ values account only for the non-bonded interaction between the two monomeric subunits. Only those hydrogen bonds that link the subunits are counted.

| Dimer | $-E_{tot}$ / kcal mol$^{-1}$ | $R_g$ / Å | $\overline{RMSD}$ / Å | $\overline{RMSF}$ / Å | $A_{IS}$ / Å$^2$ | Hydrogen bonds | Salt bridges |
|---|---|---|---|---|---|---|---|
| $cov^{189A}$ | 533 ± 122 | 32.8 ± 0.3 | 2.2 ± 0.5 | 1.4 ± 0.5 | 1407 ± 189 | 45 | 11 |

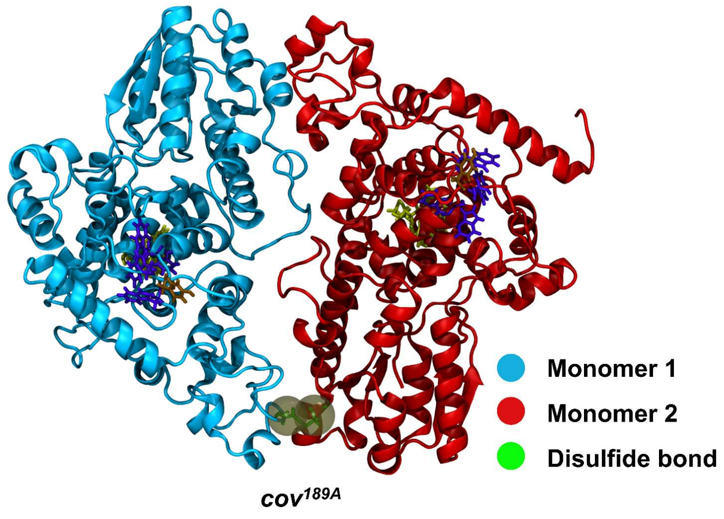


**Figure S9**: Structure of the *Er*Cry4a $cov^{189A}$ dimer. The FAD cofactor, Trp-tetrad (W395, W372, W318, W369), and Y319 are shown in yellow, violet and ochre, respectively. Brown spheres indicate the approximate position of the C189-C458 disulphide bond between the monomers.

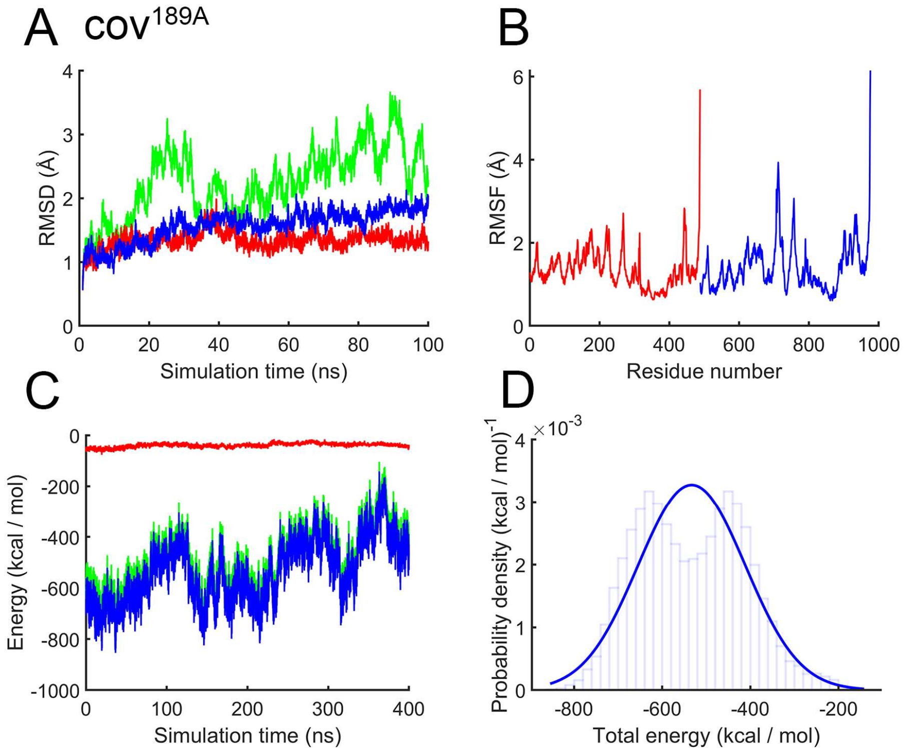


**Figure S10**: Time evolution of the root mean square deviation (RMSD) of the backbone atoms of the *Er*Cry4a cov$^{189A}$ dimer, computed relative to the structure of the corresponding dimer after equilibration (**A**). Average root mean square fluctuations $\overline{RMSF}$ values computed for the residues of the cov$^{189A}$ dimer (**B**). Red corresponds to monomer A and blue to monomer B. (**C**-**D**): Time evolution of the interaction energies between two monomeric subunits of the cov$^{189A}$ *Er*Cry4a dimer and the corresponding probability density distribution of the interaction energy. Red and green denote respectively the van der Waals and Coulomb contributions to the total interaction energy between the monomers, shown in blue.

#### *Er*Cry4a 412 dimer family

Residue C412 was also investigated computationally as a possible linker of covalent *Er*Cry4a dimers (Table S17). Eight structures were created: cov(412)$^{1,2,3,A,B,6\text{-}8}$.

The structures of the most stable dimers, cov$^{412A}$ and cov$^{412B}$ (Fig. S11), display a symmetric orientation of monomeric subunits relative to one another. Table S9 shows their $\overline{RMSD}$ values: cov$^{412A}$ and cov(412)$^{7}$ appear to be the most stable. cov$^{412A}$ has the smallest $\overline{RMSF}$ (see Fig. S13). The interaction energies (Table S9 and Fig. S14) indicate that cov$^{412A}$ and cov$^{412B}$ should be the most stable of the dimers, even though cov$^{412B}$ would be considered highly unstable based on the $\overline{RMSD}$ analysis (see Fig. S12). The strongest interaction energy (for cov$^{412A}$) is accompanied by the largest interaction surface area, suggesting that cov$^{412A}$ is the most stable dimer from the 412 family.

**Table S9**: Summary of the analysis of the cov(412)$^{n}$ dimer family similar to the results shown in Tables S4, S6, S7, S8. Interaction energy values shown here account only for non-bonded interaction between the two monomeric subunits. Only those hydrogen bonds appearing between two subunits are counted. $A_{IS}$ is the interaction surface area between the two monomeric subunits.

| Dimer | $-E_{tot}$ / kcal mol$^{-1}$ | $R_g$ / Å | $\overline{RMSD}$ / Å | $\overline{RMSF}$ / Å | $A_{IS}$ / Å$^2$ | Hydrogen bonds | Salt bridges |
|---|---|---|---|---|---|---|---|
| cov(412)$^{1}$ | −315 ± 86 | 36.9 ± 0.3 | 3.5 ± 1.0 | 6.0 ± 1.1 | 1570 ±257 | 64 | 1 |
| cov(412$^{)2}$ | −207 ± 97 | 38.9 ± 0.5 | 6.7 ± 2.3 | 7.8 ± 1.9 | 1201 ± 280 | 66 | 5 |
| cov(412)$^{3}$ | 386 ± 111 | 41.3 ± 0.4 | 3.9 ± 1.3 | 5.6 ± 1.2 | 1069 ± 330 | 40 | 20 |
| cov(412)$^{4}$= cov$^{412A}$ | 526 ± 83 | 41.2 ± 0.2 | 2.5 ± 0.4 | 5.7 ± 1.3 | 1840 ± 115 | 31 | 13 |
| cov(412)$^{5}$= cov$^{412B}$ | 437 ± 127 | 41.4 ± 0.7 | 6.2 ± 2.8 | 5.4 ± 1.5 | 1040±310 | 49 | 12 |
| cov(412)$^{6}$ | −87 ± 85 | 36.8 ± 0.4 | 3.1 ± 1.0 | 6.8 ± 1.2 | 1689 ± 248 | 65 | 5 |
| cov(412)$^{7}$ | −145 ± 53 | 38.1 ± 0.5 | 1.9 ± 0.4 | 7.0 ± 1.0 | 1274 ± 146 | 29 | 7 |
| cov(412)$^{8}$ | −382 ± 97 | 37.2 ± 0.4 | 2.9 ± 0.9 | 4.8 ± 0.7 | 1881 ± 371 | 75 | 4 |

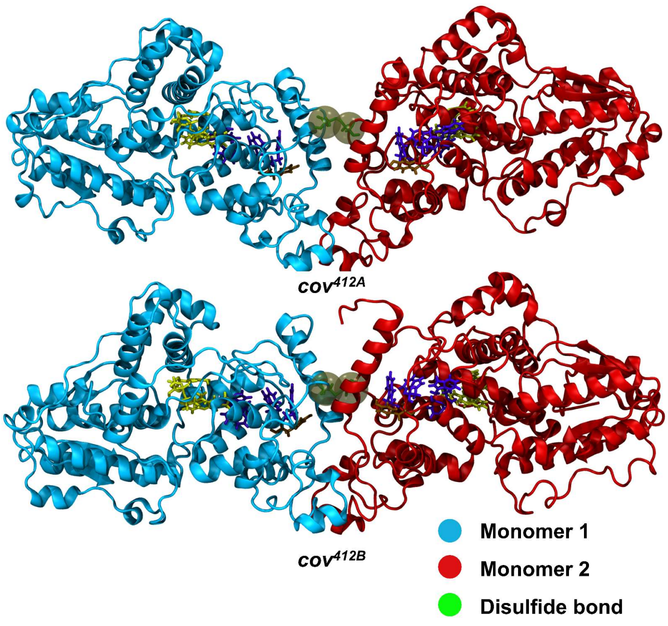


**Figure S11**: Structures of the two favourable *Er*Cry4a dimers cov$^{412A}$ and cov$^{412B}$. The FAD cofactor, Trp-tetrad (W395, W372, W318, W369), and Y319 are shown in yellow, violet and ochre, respectively. Brown spheres indicate the approximate position of the C412-C412 disulphide bond between the monomers.

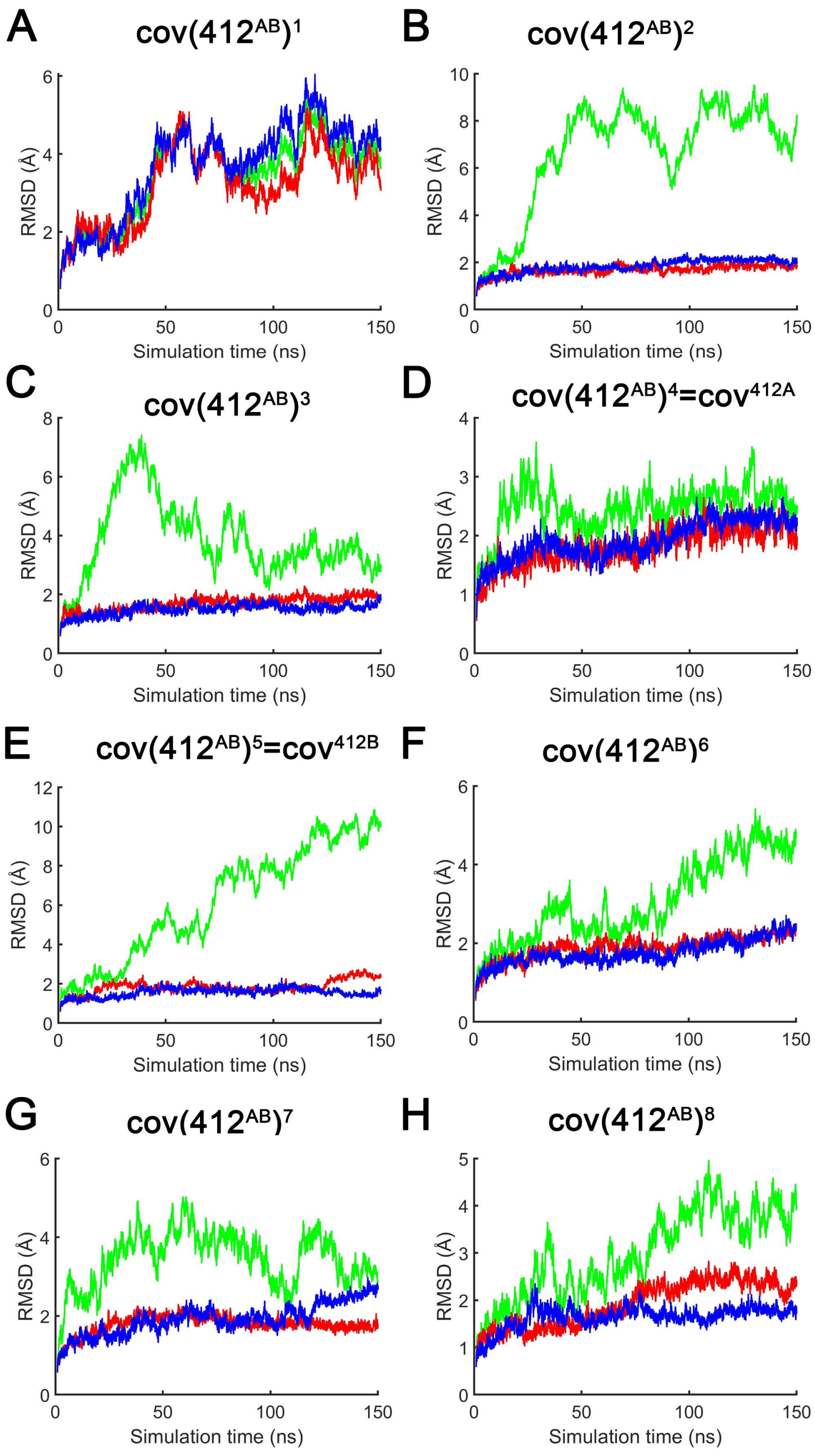


**Figure S12**: Time-evolution of the RMSD values for the *Er*Cry4a dimeric structures of the cov(412)$^n$ family, computed for each monomeric subunit (blue and red), and the whole dimer (green) relatively to the structures obtained after equilibration simulations.

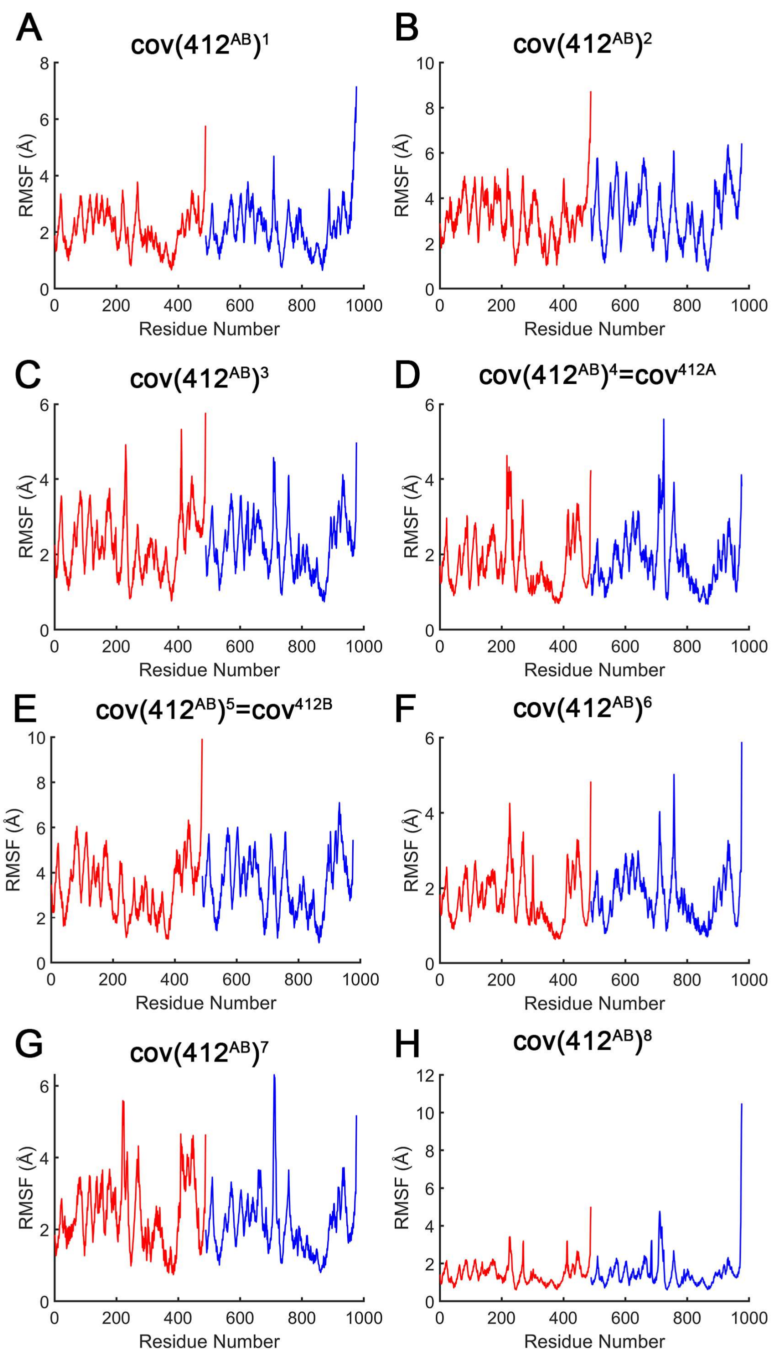


**Figure S13**: Average root mean square fluctuations $\overline{RMSF}$ values computed for the residues of the *Er*Cry4a dimeric structures of the cov(412)$^n$ family. Red corresponds to monomer A and blue to monomer B.

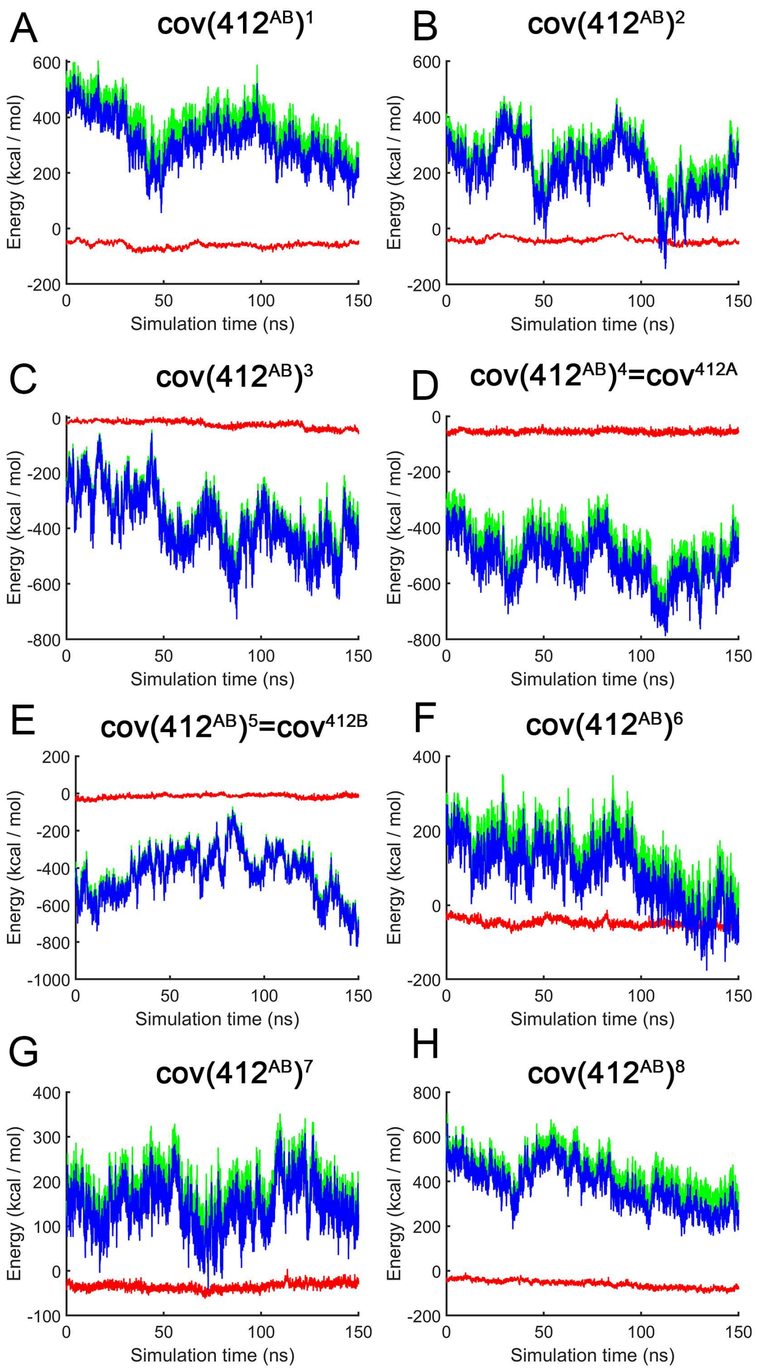


**Figure S14**: Time evolution of the interaction energies between two monomeric subunits of the cov(412)$^{n}$ *Er*Cry4a dimers. Red and green denote respectively the van der Waals and Coulomb contributions to the total interaction energy between the monomers, shown in blue.

#### Distance analysis between cysteine residues

To determine which cysteine residues are in close contact, the six most surface-exposed cysteines (C317, C116, C189, C68, C412, C73) and C179 (see Table S3) were analysed in the 49 dimeric structures obtained using the docking tools. Tables S11-S17 summarise the distances between these cysteine residues. The tables are organised by the contact residue used in the docking procedure to produce the various families of structures. Distances between cysteines less than 10 Å are coloured green, and those between 10 Å and 20 Å are in yellow. Tables S18 and S19 summarise the distances between cysteines for complexes that were docked without defining a contact residue, in one case using ZDOCK[6,7], and in the other using M-ZDOCK[5] which was used for symmetric multimer docking. The names in the following tables (Complex *N*) are used internally only here and not in the rest of this paper.

**Table S11:** Analysis of inter-monomer distances between cysteine residues in *Er*Cry4a for the 10 complexes produced by ZDOCK, where Cys68 was used as a contact residue. The distance measures the separation of the sulphur atoms in the two cysteine residues.

| ZDOCK Contact residue Cys68 | | | |
|---|---|---|---|
| | Residue 1 | Residue 2 | Distance (Å) |
| Complex 1 | 361 | 361 | 31.93 |
| | 412 | 412 | 25.56 |
| | 313 | 313 | 44.3 |
| | 257 | 257 | 56.63 |
| Complex 2 | 257 | 412 | 28.15 |
| | 179 | 458 | 44.66 |
| | 412 | 116 | 43.91 |
| Complex 3 | 257 | 458 | 23.85 |
| | 257 | 458 | 23.85 |
| | 68 | 458 | 27.79 |
| | 257 | 412 | 29.03 |
| Complex 4 | 361 | 412 | 37.72 |
| | 412 | 412 | 36.51 |
| | 257 | 257 | 41.62 |
| Complex 5 | 412 | 412 | 35.96 |
| | 458 | 458 | 72.08 |
| | 257 | 257 | 43.01 |
| Complex 6 | 412 | 412 | 14.23 |
| | 361 | 361 | 27.73 |
| | 317 | 317 | 38.97 |
| | 458 | 458 | 38.12 |
| Complex 7 | 257 | 412 | 37.24 |
| | 361 | 313 | 38.98 |
| | 313 | 313 | 41.65 |
| Complex 8 | 257 | 313 | 37.25 |
| | 68 | 412 | 36.98 |
| Complex 9 | 313 | 313 | 17.1 |
| | 361 | 361 | 30.65 |
| | 317 | 317 | 35.04 |
| | 412 | 412 | 37.86 |
| | 116 | 116 | 36.98 |
| Complex 10 | 116 | 458 | 41.95 |
| | 313 | 412 | 29.88 |
| | 317 | 361 | 36.43 |

**Table S12:** Analysis of inter-monomer distances between cysteine residues in *Er*Cry4a for the 10 complexes produced by ZDOCK, where Cys73 was used as a contact residue. The distance measures the separation of the sulphur atoms in the two cysteine residues.

| **ZDOCK Contact residue Cys73** | | | |
|---|---|---|---|
| | Residue 1 | Residue 2 | Distance (Å) |
| Complex 1 | 317 | 412 | 37.77 |
| | 412 | 412 | 26.1 |
| | 361 | 361 | 30.52 |
| Complex 2 | 412 | 257 | 29.53 |
| | 257 | 313 | 52.5 |
| | 361 | 116 | 45.41 |
| | 458 | 179 | 45.94 |
| Complex 3 | 458 | 257 | 26.98 |
| | 257 | 458 | 27.13 |
| | 458 | 68 | 26.77 |
| | 257 | 412 | 30.63 |
| Complex 4 | 313 | 458 | 46.37 |
| | 412 | 412 | 36.073 |
| | 361 | 412 | 36.023 |
| Complex 5 | 412 | 361 | 39.61 |
| | 257 | 257 | 41.84 |
| | 361 | 361 | 42.1 |
| Complex 6 | 412 | 412 | 15 |
| | 361 | 361 | 27.72 |
| | 458 | 458 | 38.13 |
| | 317 | 317 | 41.05 |
| Complex 7 | 412 | 257 | 35.24 |
| | 313 | 313 | 40.77 |
| | 361 | 116 | 47.64 |
| Complex 8 | 412 | 257 | 33.5 |
| | 412 | 179 | 34.87 |
| | 257 | 313 | 37.48 |
| Complex 9 | 313 | 313 | 19 |
| | 317 | 317 | 33.78 |
| | 361 | 361 | 34.69 |
| Complex 10 | 412 | 313 | 28.14 |
| | 361 | 313 | 28.78 |
| | 313 | 313 | 37.87 |

**Table S13:** Analysis of inter-monomer distances between cysteine residues in *Er*Cry4a for the 10 complexes produced by ZDOCK, where Cys116 was used as a contact residue. The distance measures the separation of the sulphur atoms in the two cysteine residues.

| ZDOCK Contact residue Cys116 | | | |
|---|---|---|---|
| | Residue 1 | Residue 2 | Distance (Å) |
| Complex 1 | 361 | 361 | 29.15 |
| | 412 | 412 | 27.25 |
| | 313 | 313 | 44.3 |
| Complex 2 | 179 | 458 | 45.1 |
| | 257 | 412 | 28.45 |
| Complex 3 | 458 | 257 | 24.66 |
| | 412 | 68 | 41.13 |
| | 257 | 457 | 26.61 |
| | 68 | 412 | 39.31 |
| Complex 4 | 361 | 412 | 35.42 |
| | 458 | 313 | 46.65 |
| | 68 | 68 | 40.56 |
| Complex 5 | 412 | 412 | 38.18 |
| | 412 | 361 | 41.23 |
| | 68 | 68 | 44.38 |
| | 257 | 257 | 43.01 |
| Complex 6 | 412 | 412 | 17.2 |
| | 361 | 361 | 28.26 |
| | 317 | 313 | 37.88 |
| | 458 | 257 | 30.57 |
| | 317 | 317 | 38.56 |
| Complex 7 | 257 | 412 | 34.83 |
| | 313 | 412 | 38.87 |
| | 313 | 313 | 38.78 |
| Complex 8 | 257 | 313 | 37.48 |
| | 257 | 412 | 32.32 |
| Complex 9 | 313 | 313 | 16.81 |
| | 361 | 361 | 30.56 |
| | 317 | 317 | 33.78 |
| Complex 10 | 313 | 412 | 26.58 |
| | 313 | 316 | 29.44 |
| | 313 | 313 | 39.24 |
| | 116 | 458 | 40.52 |

**Table S14:** Analysis of inter-monomer distances between cysteine residues in *Er*Cry4a for the 10 complexes produced by ZDOCK, where Cys179 was used as a contact residue. The distance measures the separation of the sulphur atoms in the two cysteine residues.

| **ZDOCK Contact residue Cys179** | | | |
|---|---|---|---|
| | Residue 1 | Residue 2 | Distance (Å) |
| Complex 1 | 412 | 412 | 26.06 |
| | 361 | 361 | 29.15 |
| | 313 | 313 | 45.8 |
| Complex 2 | 412 | 257 | 29.83 |
| | 361 | 257 | 37.23 |
| | 458 | 179 | 45.1 |
| | 361 | 116 | 45.41 |
| Complex 3 | 458 | 257 | 24.66 |
| | 257 | 458 | 24.77 |
| | 257 | 412 | 29.7 |
| | 412 | 257 | 31.31 |
| Complex 4 | 257 | 257 | 42.02 |
| | 68 | 68 | 43.15 |
| | 361 | 361 | 45.79 |
| Complex 5 | 412 | 412 | 36.98 |
| | 257 | 257 | 43.96 |
| | 361 | 412 | 42.6 |
| Complex 6 | 412 | 412 | 17.18 |
| | 361 | 361 | 27.72 |
| | 458 | 257 | 34.18 |
| | 361 | 313 | 40.26 |
| Complex 7 | 257 | 412 | 35.24 |
| | 313 | 313 | 39.28 |
| | 116 | 361 | 49.43 |
| Complex 8 | 412 | 257 | 30.51 |
| | 361 | 179 | 38.14 |
| | 257 | 116 | 42.96 |
| Complex 9 | 313 | 313 | 16.03 |
| | 317 | 317 | 35.6 |
| | 361 | 361 | 30.65 |
| Complex 10 | 412 | 116 | 30.34 |
| | 361 | 313 | 29.83 |
| | 313 | 317 | 40.48 |

**Table S15:** Analysis of inter-monomer distances between cysteine residues in *Er*Cry4a for the 3 complexes produced by ZDOCK, where Cys189 was used as a contact residue. The distance measures the separation of the sulphur atoms in the two cysteine residues.

| ZDOCK Contact residue Cys189 | | | |
|---|---|---|---|
| | Residue 1 | Residue 2 | Distance (Å) |
| Complex 1 | 189 | 189 | 15.96 |
| | 68 | 189 | 20.11 |
| | 189 | 73 | 18.96 |
| | 189 | 68 | 18.14 |
| | 189 | 257 | 18.2 |
| | 116 | 458 | 32.46 |
| Complex 2 | 189 | 189 | 15.88 |
| | 73 | 68 | 25.44 |
| | 68 | 73 | 25.58 |
| | 116 | 116 | 44.36 |
| | 116 | 257 | 35.81 |
| Complex 3 | 189 | 458 | 8.6 |
| | 189 | 458 | 25.38 |
| | 257 | 257 | 28.55 |
| | 458 | 73 | 14.4 |

**Table S16:** Analysis of inter-monomer distances between cysteine residues in *Er*Cry4a for the 3 complexes produced by ZDOCK, where Cys317 was used as a contact residue. The distance measures the separation of the sulphur atoms in the two cysteine residues.

| **ZDOCK Contact residue Cys317** | | | |
|---|---|---|---|
| | Residue 1 | Residue 2 | Distance (Å) |
| Complex 1 | 317 | 317 | 4.39 |
| | 313 | 313 | 20.03 |
| | 361 | 361 | 29.72 |
| | 317 | 313 | 14.66 |
| | 313 | 317 | 13.78 |
| Complex 2 | 313 | 313 | 18.12 |
| | 317 | 317 | 21.93 |
| | 317 | 313 | 21.19 |
| | 313 | 317 | 21.68 |
| | 361 | 317 | 28.45 |
| Complex 3 | 317 | 116 | 4.11 |
| | 458 | 313 | 16.01 |
| | 458 | 317 | 13.67 |
| | 412 | 313 | 24.08 |

**Table S17:** Analysis of inter-monomer distances between cysteine residues in *Er*Cry4a for the 3 complexes produced by ZDOCK, where Cys412 was used as a contact residue. The distance measures the separation of the sulphur atoms in the two cysteine residues.

| **ZDOCK Contact residue Cys412** | | | |
|---|---|---|---|
| | Residue 1 | Residue 2 | Distance (Å) |
| Complex 1 | 412 | 412 | 31.74 |
| | 361 | 361 | 37.26 |
| | 313 | 313 | 40.47 |
| Complex 2 | 458 | 458 | 17.72 |
| | 412 | 412 | 18.15 |
| | 361 | 421 | 21.83 |
| | 317 | 361 | 33.03 |
| Complex 3 | 313 | 412 | 23.46 |
| | 361 | 361 | 23.01 |
| | 412 | 412 | 21.23 |
| | 412 | 317 | 23.9 |

**Table S18:** Analysis of inter-monomer distances between cysteine residues in *Er*Cry4a for the 10 complexes produced by ZDOCK, where no contact residue was specified. The distance measures the separation of the sulphur atoms in the two cysteine residues.

| **ZDOCK NO Contact residue** | | | |
|---|---|---|---|
| | Residue 1 | Residue 2 | Distance (Å) |
| Complex 1 | 68 | 68 | 33.21 |
| | 257 | 257 | 40.03 |
| | 179 | 179 | 34.49 |
| Complex 2 | 257 | 257 | 65.64 |
| | 412 | 412 | 55.1 |
| | 68 | 68 | 64.26 |
| Complex 3 | 412 | 68 | 36.84 |
| | 458 | 68 | 37.34 |
| | 361 | 257 | 39.36 |
| Complex 4 | 313 | 412 | 35.31 |
| | 257 | 257 | 41 |
| | 68 | 68 | 48.34 |
| Complex 5 | 412 | 412 | 19.23 |
| | 361 | 68 | 15.65 |
| | 313 | 73 | 26.53 |
| | 317 | 189 | 29.74 |
| Complex 6 | 257 | 257 | 43.35 |
| | 68 | 179 | 41.68 |
| | 179 | 68 | 39.77 |
| Complex 7 | 257 | 412 | 39.18 |
| | 257 | 361 | 41.8 |
| | 361 | 313 | 39.63 |
| Complex 8 | 458 | 317 | 22.01 |
| | 257 | 361 | 33.16 |
| | 68 | 412 | 28.62 |
| | 458 | 313 | 13.12 |
| Complex 9 | 412 | 313 | 31.13 |
| | 361 | 313 | 32.41 |
| | 313 | 313 | 37.34 |
| Complex 10 | 317 | 317 | 35.95 |
| | 313 | 361 | 29.48 |
| | 313 | 412 | 26.85 |
| | 313 | 313 | 33.94 |
| | 361 | 313 | 34.27 |

**Table S19:** Analysis of inter-monomer distances between cysteine residues in *Er*Cry4a for the 10 complexes produced by M-ZDOCK, where no contact residue was specified. The distance measures the separation of the sulphur atoms in the two cysteine residues.

| **M-ZDOCK NO Contact residue** | | | |
|---|---|---|---|
| | Residue 1 | Residue 2 | Distance (Å) |
| Complex 1 | 189 | 189 | 61.87 |
| | 73 | 189 | 20.28 |
| | 189 | 73 | 20.35 |
| Complex 2 | 313 | 116 | 21.87 |
| | 116 | 313 | 19.44 |
| | 116 | 317 | 19.69 |
| | 116 | 361 | 21.34 |
| Complex 3 | 412 | 412 | 30.33 |
| | 361 | 361 | 31.3 |
| | 313 | 313 | 31.76 |
| Complex 4 | 313 | 116 | 16.8 |
| | 317 | 116 | 20.71 |
| | 116 | 313 | 19.39 |
| | 116 | 317 | 19.7 |
| Complex 5 | 313 | 116 | 19.74 |
| | 317 | 116 | 20.35 |
| | 116 | 313 | 19.74 |
| | 116 | 317 | 19.064 |
| Complex 6 | 313 | 313 | 30.59 |
| | 257 | 361 | 42.41 |
| Complex 7 | 189 | 189 | 30.59 |
| | 458 | 458 | 41.37 |
| | 458 | 458 | 30.92 |
| | 458 | 412 | 32.3 |
| Complex 8 | 116 | 313 | 19.87 |
| | 116 | 317 | 20.45 |
| | 313 | 116 | 18.454 |
| | 116 | 361 | 20.915 |
| Complex 9 | 116 | 317 | 5.09 |
| | 116 | 313 | 6.02 |
| | 313 | 116 | 8 |
| | 317 | 116 | 2.2 |
| Complex 10 | 313 | 313 | 20.34 |
| | 116 | 313 | 21.54 |
| | 317 | 116 | 23.46 |

#### Molecular dynamics simulation protocol

**Table S20:** Summary of all the MD simulations performed. cov and ncov denote covalent and non-covalent dimers, respectively. aa stands for amino acid. cov$^{D}$ is the dimer with two disulphide bonds (Cys116-Cys313 and Cys313-Cys116). ncov$^{M}$ is the non-covalent dimer based on the structure of mouse Cry2.

| Dimer | Statistical ensemble | Integration time step (fs) | Constrained atoms | Simulation time (ns) |
|---|---|---|---|---|
| **Initial equilibration** | | | | |
| (cov317)$^{A,B,3}$ | NPT | 1 | Protein except aa 310-320 | 1 |
| | NPT | 1 | Backbone except 310-320 | 2 |
| | NVT | 1 | None | 2 |
| (cov412)$^{1,2,3,A,B,6-8}$ | NPT | 0.1 or 1 | Protein except aa 410-414 | 1 |
| | NPT | 1 | Backbone except 410-414 | 15 |
| | NVT | 1 | None | 15 |
| (cov189) | NPT | 1 | Protein | 1 |
| | NPT | 1 | Backbone | 2 |
| | NVT | 1 | None | 3 |
| cov(116$^{A}$313$^{B}$-313$^{A}$116$^{B}$)= cov$^{D}$ | NPT | 1 | Protein | 1 |
| | NPT | 1 | Backbone | 5 |
| | NVT | 1 | None | 5 |
| mouse-like = ncov$^{M}$ | NPT | 1 | Protein | 1 |
| | NPT | 1 | Backbone | 5 |
| | NVT | 1 | None | 5 |
| ncov$^{1-3}$,ncov$^{A,5-9}$ | NPT | 2 | None | 2 |
| **Production simulation** | | | | |
| (cov317)$^{1-3}$/ | NVT | 2 | None | 400 |
| (cov412)$^{1,2,3,A,B,6-8}$ | NVT | 2 | None | 150 |
| (cov189) | NVT | 2 | None | 100 |
| cov$^{D}$ | NVT | 2 | None | 100 |
| ncov$^{M}$ | NVT | 2 | None | 100 |
